# Lineage of origin-specific developmental programs drive the behaviors of malignant cells in an avian embryo model of human Medulloblastoma subgroups

**DOI:** 10.1101/2025.10.09.681440

**Authors:** Marion Mallet, Florian Martin, Karine Thoinet, Caroline Imbert, Moein Sarhadi, Jeremy Ganofsky, Jacob Torrejon Diaz, Tanguy Fenouil, Céline Delloye-Bourgeois, Julien Falk, Cécile Faure-Conter, Ludovic Telley, David Meyronet, Olivier Ayrault, Valérie Castellani, Servane Tauszig-Delamasure

## Abstract

Tumoral cells of medulloblastoma (MB) subgroups (SHH, G3 and G4) display a close transcriptomic proximity to early neuronal progenitors that migrate to form the embryonic cerebellum. With the aim of exploring functional proximities between MB cells and their physiological counterparts, we established a model of transplantation of human MB cells into the cerebellum of chick embryos in vivo and ex ovo. Light-sheet imaging of embryos grafted with cell lines and patient biopsies of MB SHH, G3 and G4 revealed the formation of primary tumors within a few days, whose topography matched that of the different MB subgroups on patients’ MRI. We found that transplanted MB cells adopted morphological and migratory features specific of their respective lineage of origin. Combining transcriptomic and functional approaches, we found that MB G3 tumoral cells exploit the canonical SLIT migration developmental signaling at disease emergence. This signaling is maintained in MB G3 patients and is associated with tumor aggressivity and poor prognosis.

## INTRODUCTION

Medulloblastoma (MB) is a malignant embryonic tumor of the developing cerebellum. It is the most common pediatric malignant brain tumor. The standard treatments relying on surgical resection, radiation and chemotherapy have led to favorable short-term outcomes overall, but are often associated with long-term sequelae in children^1^. MB has been classified into four main subgroups (WNT, SHH, Group 3 (G3), and Group 4 (G4)) with different etiologies, presentations and prognoses. The animal models currently used to study MB rely mostly on genetically engineered or immunodeficient adult mice grafted with patient-derived samples (PDX) (for review^2^). Strikingly, there is no animal model for MB G4 compatible with research so far. This issue might be due to complexity in modeling the embryonic context in which this cancer emerges.

Indeed, MB result from alterations occurring during early embryonic cerebellum development. Recent studies comparing the single-cell transcriptome of MB tumors with transcriptomic atlases of human early cerebellar progenitors revealed a close proximity of tumoral cells with different progenitors derived from the rhombic lip (RL). The RL germinal zone indeed gives rise to the progenitors of the cerebellar glutamatergic excitatory neurons: the Cerebellar Nuclei (CN), the Granular Cells (GC) and the Unipolar Brush Cells (UBC)^3–5^ (Fig. 1a).

**Figure 1.**
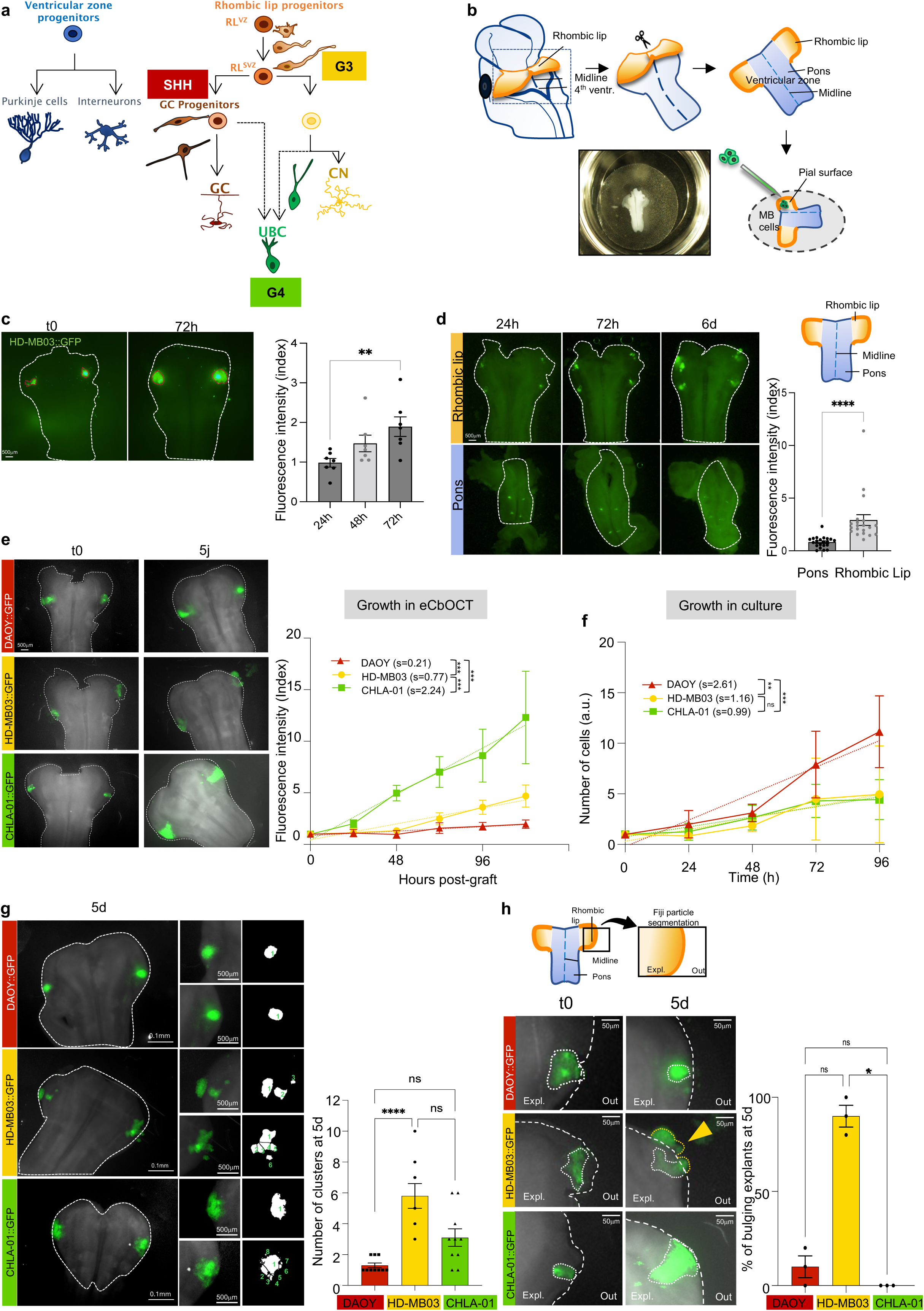
The graft of MB cells in whole tissue culture of the chick primitive cerebellum (embryonic Cerebellar Organotypic Tissue Culture, eCbOTC) allows the observation and monitoring of tumor mass formation and reveals subgroups differences. **a.** Lineage trees of early cerebellar progenitors, lineages of origin of MB SHH, G3 and G4. Progenitors emerge from the two cerebellar germinal zones: (i) the ventricular zone (VZ) gives rise to the future inhibitory GABAergic neurons (Purkinje cells) and interneurons. (ii) The RL gives rise to the progenitors of excitatory Glutamatergic neurons: excitatory Cerebellar Nuclei progenitors (CN), Granular Cell Progenitors (GCP) and Unipolar Brush Cells (UBC) progenitors. In humans, the RL is compartmented; progenitors transit from the RL lining the 4^th^ ventricle (RL^VZ^) to the RL subventricular zone (RL^SVZ^) to undergo differentiation. MB SHH originate from GC, MB G3 from early RL and CN and MB G4 from UBC respective lineages. The lineage of UBC is still debated, they might have a common ancestor with CN or derive from GC progenitors (dotted arrows). **b.** Schematic representation of eCbOTC experimental set-up. A representative picture is shown on the inset of the “open-book” explant of the RL and pons spread on a membrane and floating on the medium. c. Monitoring of tumor mass formation by HD-MB03::GFP cells grafted in eCbOTc. Representative pictures of t0 and 72h grafted eCbOTC are shown on the left. Fluorescence intensity of the tumor mass was measured (dotted red line) just after the graft (t0), 24h, 48h and 72h post-graft and a ratio with the t0 value was calculated. The Mann-Whitney test was used to compare each condition with 24h control condition. **d.** Comparison of the tumor mass growth in the RL versus in the adjacent pons, 6 days (6d) post-graft. The Mann-Whitney test was performed between both conditions at 72h as at 6d some cells bulge out of the explant (see Fig. 1h), biasing the quantification. Representative pictures are shown on the left. Dotted white lines delineate the explant including the rhombic lip (upper panel) and the pons only (lower panel). **ef.** Comparison of the rate of tumor mass growth in eCbOTC (e) versus in culture (f) for MB SHH (DAOY::GFP in red), MB G3 (HD-MB03::GFP in yellow) and MB G4 cells (CHLA-01::GFP in green). Dotted lines on the diagram represent the regression of the growth curves. The corresponding slopes (s) are indicated. Representative pictures are presented on the left with white dotted lines delimiting the explants. Equivalent of ANCOVA test was performed to compare the slopes by pairs. **g.** Macroscopic observations of more or less cohesive tumor masses formed by cells lines corresponding to the different MB subgroups. Clusters of cells were quantified for tumors masses formed by DAOY::GFP, HD-MB03::GFP and CHLA-01::GFP, 5 days (5d) post-graft. Representative pictures are shown on the left with zoomed inserts with the quantification of cell clusters indicated in green. White dotted lines delimit the explants. The Kruskal-Wallis test was performed to compare the mean number of clusters for each cell line. h. Observation and monitoring of tumor mass bulging out of the eCbOTC, 5 days (5d) post-graft. Schematic representation on the upper inset and white dotted lines in the pictures delimitate what is considered in and out of the explant. The striking bulging of MB G3 is indicated by a yellow arrowhead. The Kruskal-Wallis test compares the mean percentage of explants with observed bulging at 5 days post-graft. Error bars indicate SEM. ns non-significant, * p≤0.05, ** p≤0.01, *** p≤0.001, **** p≤0.0001.

In Humans, the RL germinal zone is compartmentalized in two anatomically and transcriptionally distinct regions: the ventricular RL (RL^VZ^) generates the earliest progenitors which then transit to the RL^SVZ^ (RL subventricular zone) to start their differentiation^6,7^. The compartmentation of the germinal zone seem to be slightly different in mice and chicken, yet, in these 3 species, cerebellar neuronal progenitors emerge in temporal waves from the RL caudal edge: first the CN progenitors (around 7-17PCW in Human, E10.5-E12.5 in mice, E3 in chick), second, the GC progenitors (GCP) (around 10-18PCW in Human, E12.5 in mice, E4 in chick), and later the UBC progenitors (around 16-25 PCW in Human, E14 in mice, E6 in chick)^8–11^ (Suppl. Fig. 1a). These progenitors proliferate and commit into their differentiation lineage while they migrate toward their final location in the different layers of the stratified cerebellar neuroepithelium (Suppl. Fig. 1a). When emerging from the edge of the RL, these progenitors already display morphologic specificities and stereotyped migration modes and trajectories. CN and GCP initially switch from a round amoeboid to a unipolar shape, they extend a leading process and migrate both tangentially along the pial surface of the RL. CN then migrate the most rostrally toward the Nuclear Transitory Zone (NTZ) and later radially toward the future white matter to coalesce and form the Deep Cerebellar Nuclei (DCN)^10,12^ (Suppl. Fig. 1a). GCP stop at the pial surface and proliferate massively to form the External Granular Layer (EGL). They adopt a bipolar shape and will later extend a third perpendicular axon to migrate radially toward the Inner Granular Layer (IGL). Finally, UBC progenitors migrate more ventrally through the nascent white matter to eventually differentiate and establish synapses connecting GC to mossy fibers in the IGL at the border of the white matter. UBC are particularly enriched in the flocculonodular lobes of the caudal cerebellum, that line the 4^th^ ventricle ^10,13,14^(Suppl. Fig. 1a).

The studies comparing MB tumors with human embryonic transcriptomic atlases have confirmed the putative cells of origin of MB WNT among a population of extracerebellar progenitors. For MB SHH, they confirmed the GCP as the cells of origin, consistent with previous mouse modeling. They also stated that MB G3/G4 would constitute a continuum of more or less differentiated tumors, along the same physiological lineage, distinct from that of the GCP, including RL^VZ^, early RL^SVZ^, CN and early UBC progenitors which would give rise to MB G3 while MB G4 would derive from more committed RL^SVZ^ and UBC progenitors^3–6^. However, this hypothesis still needs to be tested experimentally, as a recent study using a different dataset suggests that MB G3/G4 may derive from an alternative GCP-UBC lineage^15^ (Fig. 1a). Lastly, it is still unknown whether this transcriptomic proximity of tumor cells with these neuronal lineages confers functional specificities to MB cells.

We hypothesized that replacing MB cells back into their embryonic territory of origin might reactivate these lineage-dependent features and could enlighten developmental mechanisms that tumoral cells might exploit during MB onset. By grafting human MB tumoral cells into the RL of chick embryos we succeeded to engineer the formation of tumors within a few days for all MB subgroups, including MB G4. In our model, the topography of the tumors observed in the chick embryos by light-sheet microscopy was comparable to that observed in patients by MRI. At the microscopic level, we observed that MB cells, once grafted into the primitive cerebellum, acquired morphologies fully reminiscent of their lineage of origin. Finally, we compared the transcriptome of MB G3 cells, before and once settled into the chick primitive cerebellum. This analysis revealed that MB cells experiencing the microenvironment of their territory of origin activate transcriptomic programs associated with a proliferative undifferentiated state that match those of MB patients. In addition, we found prominent activation of SLIT signaling in grafted cells. MB G3 express the SLIT receptors ROBO1 and PLXNA1 and SLIT2 is expressed early during cerebellar development at the edge of the RL. Using our avian model and ex vivo cultures, we found that SLIT2 loosens MB cell-cell cohesion while increasing the exploratory and infiltrative properties of the forming primary tumor. Thus, MB cells adapt to the embryonic microenvironment and take advantage of local signals to promote their malignant behaviors.

## RESULTS

### Early MB tumor formation can be modelized in chick embryonic cerebellum organotypic tissue culture

With the aim of replacing MB cells into the specific embryonic context in which this cancer emerges, we set up the graft of MB cell lines into organotypic tissue culture of chick embryo primitive cerebellum (embryonic Cerebellum Organotypic Tissue Culture, eCbOTC). The RL and pons of HH26-E5 (Hamburger-Hamilton stage 26, embryonic day 5) chick embryos were dissected. “Open book” preparations were achieved by incision along the roof plate and placed on a microporous membrane floating at the surface of the culture medium (Fig. 1b and Material and Methods). eCbOTC were cultured for at least 5 days. During this culture period, tissues survive and explants grow significantly in size, over the first 48h (Supp. Fig. 1b). Although the 3D cerebellar development is partially altered by the disruption of the roof plate, the early cerebellar cytoarchitecture is conserved: SOX2+ progenitors localize at the edge of the RL in the expected germinal zone and are in close proximity with the TUJ1-positive axonal network formed by GCP progenitors (Supp. Fig. 1c). Different MB cell lines were selected in each of the MB subgroups based on the classification reported by Ivanov and colleagues^16^ : DAOY cells for MB SHH^17^, HD-MB03 for MB G3^18^ and CHLA-01-MED (CHLA-01 for convenience) for MB G4^19^. We generated stable cell lines expressing GFP through lentiviral infection and grafted them onto the pial surface of the eCbOTC as close as possible to the edge of the RL. We first found that grafted HD-MB03::GFP (HD-MB03 for simplicity) cells, survive, aggregate and form tumor masses whose fluorescence intensity increases significantly over the first 72h of culture, with a reproducible rate among the same batch of explants (Fig. 1c). To determine if MB cells take advantage of the co-cultivated tissue, we compared their survival when deposited as patches directly on the membrane. We found that under this condition, cells do not survive after 3 days (Supp. Fig. 1d). This model thus allows a quantitative monitoring of MB tumor growth within embryonic tissues (Fig. 1c). Notably, we wondered whether the RL exerts specific effects, so we compared MB tumor growth in these RL preparations and in tissue prepared from the pons, which is the adjacent - non-cerebellar - territory. Interestingly, we observed in this latter condition the formation of tumoral masses that were more than 3.5 times smaller over the same time period (72h post-graft). 6 days after the graft, tumoral masses had significantly grown in the RL and almost disappeared in the pons (Fig. 1d).

Next, to determine whether all MB cell lines are able to grow in this specific embryonic microenvironment and whether it has different impact on their growth, we grafted the 3 MB cell lines in parallel in the eCbOTC. We found that all cell lines formed growing tumoral masses over 5 days (Fig.1e). Interestingly, whereas DAOY grew more rapidly in vitro than HD-MB03 and CHLA-01, which have comparable growth rates between them (Fig. 1f), once grafted in their territory of origin, CHLA-01 (slope of regression curve (s), s=2.24) and HD-MB03 (s=0.77) grew at a faster rate than DAOY cells (s=0.21) (Fig. 1e). This suggests that the embryonic microenvironment of the primitive cerebellum has an impact on the growth of MB cells and this impact seems to be subgroup dependent. Besides different proliferative properties, we also observed cell-lines-specific characteristics. While DAOY cells formed round, plain, cohesive and well delimited masses, HD-MB03 formed heterogeneous and more dispersed cell clusters. Amidst the two subgroups, CHLA-01 exhibited an intermediate profile and seemed to colonize the territory as sheets. By quantifying the number of cell clusters identified within the mass, we confirmed the lower cohesion of CHLA-01 and HD-MB03 masses, compared to DAOY ones (Fig.1g). Interestingly, and for HD-MB03 only, we frequently observed that tumor masses were able to bulge exophytically out of the explant after 5 days in culture (Fig. 1h). This specific behavior might reflect the infiltrative capacities of the cells and the specific bulging of MB G3 into the lumen of the 4^th^ ventricle as described by Northcott and colleagues in patients’ MRI^3^.

### The graft of MB cells in the organotypic culture of chick primitive cerebellum reveals behaviors reminiscent of their lineages of origin

In order to further characterize the eCbOTC model at the cellular and molecular scale, staining of progenitors (SOX2) and neuronal projections (β-Tubulin III (TUJ1)) was performed on the whole explant, 5 days after the graft of HD-MB03. We observed that MB cells aggregated in close proximity with their physiological counterparts within the primitive cerebellum, *i.e.* the progenitors emerging from the germinal zone at the edge of the RL (Fig. 2a). At a higher resolution, we could observe that MB cells were integrated into the dense network of neuronal cellular bodies and neurites (Fig. 2b). Some MB cells remodeled, extending protrusions that come in close contact with TUJ1-positive axons, thus suggesting that they interact with the embryonic microenvironment and their surrounding physiological counterparts (Fig. 2b). These morphological and polarity changes might be instructive of the migration and navigation modalities of the MB cells during tumor onset. As MB cells display a close transcriptomic proximity with their respective lineage of origin, we investigated whether they were able to reactivate these lineage specific morphological programs. We thus established classes of morphologies that have been described for early RL derived cerebellar progenitors: early proliferative RL protrusive progenitors, unipolar progenitors of CN or GCP tawed by short (<20μm) and later long (>20μm) processes, bipolar GCP and unipolar brush UBC progenitors^20,21^ (Fig. 1a). We analyzed the morphologies of the MB subgroups-representative cell lines in culture and 5 days post-grafting in the eCbOTC. Amongst cells with identifiable morphology, we observed that DAOY::GFP that are surrounded by large lamellipodia in culture, extended long neuronal-like processes and adopted a bipolar shape (Fig. 2c); HD-MB03 which were quite round in culture mostly extended small protrusions (Fig. 2d); CHLA-01 that were round and grew in suspension in culture extended very atypical bi-ramified protrusions (Fig. 2e). We quantified these morphological features for at least 3 embryos grafted with each cell line. DAOY cells (MB SHH subgroup) were statistically more likely to adopt the bipolar shape of GCP, the MB SHH lineage of origin. HD-MB03 cells (MB G3) preferentially adopted a morphology corresponding to early RL progenitors. CHLA-01 cells (MB G4) predominantly exhibited the prototypical brush morphology of their UBC lineage of origin (Fig. 2f and Supp. Fig. 2a). These observations suggest that, when exposed to the embryonic microenvironment, MB cells reveal a functional proximity with their respective lineage of origin, while also expressing their malignant features.

**Figure 2.**
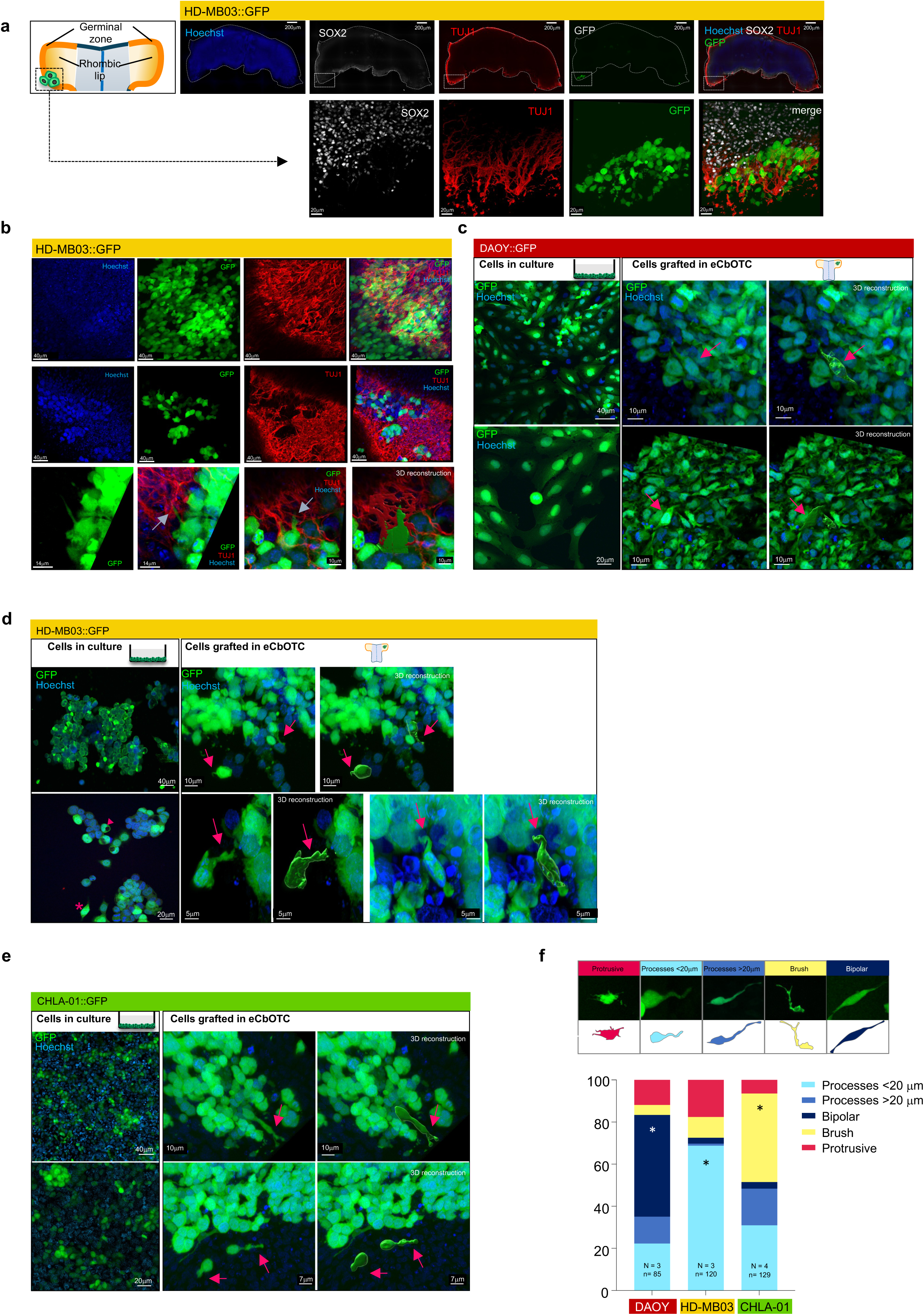
Once grafted in eCBOTC, MB cells integrate into the tissue and adopt morphologies reminiscent of their lineage of origin. **a.** Whole-tissue immunostaining of eCbOTC grafted with HD-MB03::GFP using Hoechst (nuclei), SOX2 (progenitors), β-Tubulin III (TUJ1) (neuronal projections) antibodies. Representative images acquired with Thunder optonumerical microscope are shown. White dotted lines delineate the explant (upper panel). The lower panel displays zooms corresponding to the upper panel insets. **b.** Whole tissue staining of eCbOTC grafted with HD-MB03::GFP using Hoechst, SOX2 and TUJ1 antibodies and imaged with a confocal microscope. The upper panel displays a representative image from a tumor mass embedded in neuronal projections. The middle panel displays the observation of a smaller tumor mass with peripheral cells extending processes in contact with axons (white arrow, lower panel). 3D reconstruction was performed using Imaris software to visualize the processes extended by tumoral cells. **c.** Longitudinal eCbOTC cryosections imaged with confocal microscope were performed to study cell morphology. Representative images of DAOY::GFP in culture (left panel) or 5 days post-graft in eCbOTC (right panel) are shown. The bipolar shape characteristic of GCP is indicated with a pink arrow. 3D reconstruction was performed using Imaris software to reconstruct the surface of the stained tumoral cell. **d.** Representative images of HD-MB03::GFP in culture (left panel) or 5 days post-graft in eCbOTC (right panel). In culture, HD-MB03 are predominantly round in culture (pink arrowhead) and rarely fusiform (pink asterisk). Once grafted in the primitive cerebellum, HD-MB03::GFP cells extend small processes (pink arrows). **e.** Representative images of CHLA-01::GFP in culture (left panel) or 5 days post-graft in eCbOTC (right panel). Cells harboring a brush of fine dendrioles characteristic of UBC are labelled with pink arrows. **f.** Striking morphological features (upper panel) were quantified for the different MB subgroups. They were calculated as a percentage of the total amount of cells displaying one of the morphologies indicated. The number of eCbOCT (N) and cells (n) analyzed are indicated. For each morphology, Mann-Whitney tests were performed to compare two cell lines at a time. Only statistically significant features specific of one condition are indicated. * p≤0.05.

### MB graft in the primitive cerebellum of chick embryo in vivo recapitulates MB early tumor growth with subgroup specific locations

These findings support that the xenografting of human MB cells in relevant avian embryonic tissue allows the study of their behaviors. However, the fact that the 3D cerebellum development was constrained in organotypic culture was an important limitation. So, we set up the graft of MB cells directly in the RL of living chick embryos. A significant advantage of the chick embryo model resides in the fact that, conversely to mice and human embryos, the cerebellum development is achieved at hatching^22^. In addition, birds and mammals share the evolutive acquisition of the proliferation, migration and differentiation mechanisms at the origin of cerebellar neuroepithelium stratification^22^. At E14 (HH40), the chick cerebellum displays folia and lobes with a developing vasculature corresponding approximately to 14 PCW stage in human and P6 stage in mice (Supp. Fig. 3abc)^23^. Embryos were grafted during early developmental stages at E3 (HH18) when progenitors emerge from the RL^8^ (Supp. Fig. 1a). Grafted embryos were first collected 2 to 4 days after the graft (E5-E7, HH26-31) and underwent in toto immunostaining with an anti-GFP antibody and an antibody which specifically targets a human antigen (α-mitochondria). All the grafted MB cell lines stably expressing GFP were stained with α-mitochondria antibody consistently while the host cells were not (Supp. Fig. 4a). Stained embryos were then cleared and imaged by Selective Plane Illumination Microscopy (SPIM) and analyzed in 3D (Fig. 3a, Movies S1 and S2). Autofluorescence of the tissues was used for 3D visualization of the structures (Fig. 3b). We first analyzed E5 embryos grafted at E3 with HD-MB03 and double-stained with α-GFP and α-mitochondria antibodies. In optic sections, we observed small masses of double-stained cells in the RL (Fig. 3cd). We established a topographical map of the localization of the tumor masses observed in the embryos grafted for the 3 cell lines by refining the localization of the tumor on the medio-lateral axis with coronal optic sections and on the antero-posterior axis with sagittal optic sections. We observed three preferential homing compartments: (i) RL (RL) (Fig. 3d), (ii) the cerebello-pontine angle (CPA) (Fig. 3e), and (iii) the meninges (M), where, more rarely, small masses or isolated cells were observed at the level of the roof of the 4^th^ ventricle or around the RL and the pons (Supp. Fig. 4b). Similarly in embryos grafted with DAOY (Fig. 3f) and CHLA-01 (Fig. 3g) cells, we observed tumor masses in the RL or in its close vicinity for both cell lines. To gain additional insights, we studied two other cell models: PTC1MUT murine SHH MB cells from the *Ptch1*^+/-^ mouse model^24^, and the MED8A cells representative of MB G3^16^. We found that MED8A and PTC1MUT also established tumor masses visible 2 days after the graft in the RL, the CPA or the meninges with no specific location (Supp. Fig. 4cd). Notably in the avian model, grafting hundreds of CHLA-01 cells was sufficient to drive the formation of a tumor within 2 days, while it takes more than 45 days for 2.10^5^ cells to grow in NOD/SCID mice^19^, thus indicating that MB cells rapidly adapt to the microenvironment of emergence. These topographic observations were quantified and we observed no significant difference among the MB subgroups at this early stage (Fig. 3h).

**Figure 3.**
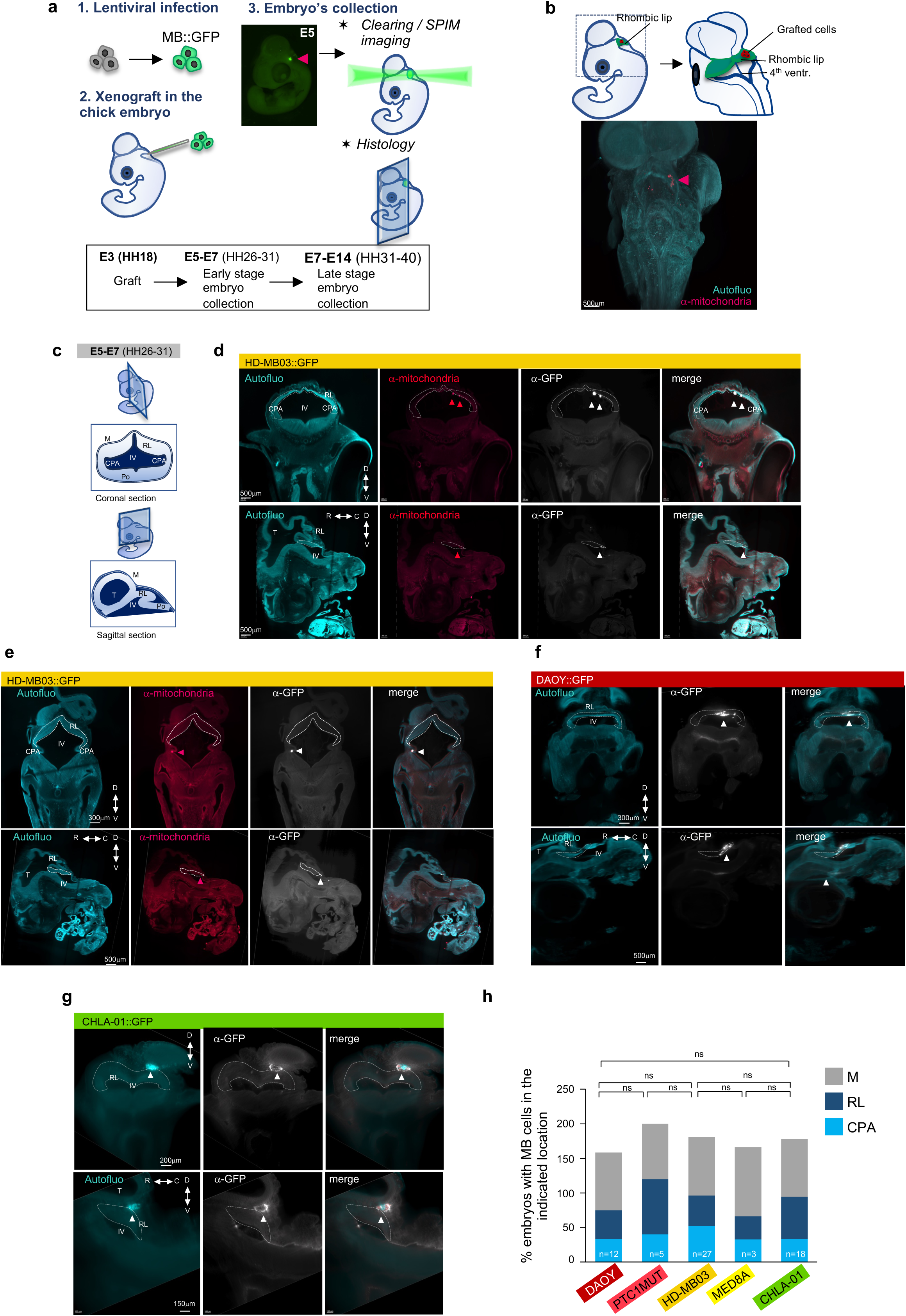
MB cells grafted in the chick cerebellum primordium aggregate to form tumoral masses. **a.** Overview of fluorescent human MB HD-MB03 cells graft paradigm in the chick embryo. A representative picture of the grafted embryo observed with a stereomicroscope is shown (right panel, arrowhead labels the tumor mass) before clearing. **b.** Representative Selective Plane Illumination Microscopy (SPIM) picture of a whole chick embryo 72hr post-graft of HD-MB03::GFP cells, cleared and labeled with anti-human mitochondria antibody. The arrowhead points to the tumoral cells. See also Movies S1 and S2. **c.** Schematic representation of coronal (upper panel) and sagittal section (lower panel) of the primitive cerebellum of a chick embryo at early stage (E5-E7, HH26-31). **d.** Representative coronal (upper panel) and sagittal (lower panel) optic sections of a chick embryo 72h post graft of HD-MB03::GFP in toto stained with an anti-mitochondria antibody targeting human cells and an anti-GFP antibody. Autofluorescence (Autofluo) allows the visualization of structures. The whole embryo was imaged with SPIM. Arrowheads point to the tumoral masses in the RL (RL) delimited by white doted lines. **e.** Representative coronal (upper panel) and sagittal (lower panel) optic sections of a chick embryo 72h post graft of HD-MB03::GFP as in d. The tumor mass is observed in the Cerebello-Pontine Angle (CPA). **f.** Representative coronal (upper panel) and sagittal (lower panel) optic sections of a chick embryo 72h post graft of DAOY::GFP in toto stained with an anti-GFP antibody. The whole embryo was imaged with SPIM. Arrowheads point to the tumoral mass. **g.** Representative optic sections of a chick embryo 72h post graft of CHLA-01::GFP as in f. **h.** Quantification of the percentage of embryos at early stages (E5-E7, HH26-31) with tumor masses in the indicated location. n is the number of embryos grafted with the indicated cell lines which were analyzed. The Chi-squared test was performed to compare two cell-lines and two locations at a time. There was no significant discrepancy for the presence of the different cell lines in the 3 locations. M: meninges, RL: RL, CPA: cerebello-pontine angle, IV: fourth ventricle, Po: Pons, T: tectum, Cb: cerebellum, D: dorsal, V: ventral, R: rostral, C: caudal, A: anterior, P: posterior, ns: non-significant.

Next, to observe MB behaviors at prolonged stages, we conducted a time course analysis from 4 to 11 days after the graft (E7-E14, HH31-40), when the chick cerebellum has expanded and formed folia (Fig. 4a). At these stages, HD-MB03 cells formed very large dispersed tumors at the border of the 4^th^ ventricle (Paraventricular localization, PV) (Fig. 4b, Movie S3), comparable to those formed by MED8A (Supp. Fig. 4e). DAOY and PTC1MUT cells formed more cohesive and plain tumors located at the surface of the vermis (Ve) or at the periphery of the cerebellar hemispheres (CH), predominantly in the caudal region of the cerebellum (Fig. 4c and Supp. Fig. 4f). Finally, CHLA-01 cells formed poorly cohesive and large paraventricular tumors (Fig. 4d). Close comparative cartography revealed that DAOY and PTC1MUT, MB SHH cells, are significantly more likely to form tumors at the periphery of the cerebellum (V/CH) whereas HD-MB03, MED8A and CHLA-01 tumors - MB G3 and G4 respectively - predominantly line the 4^th^ ventricle (Fig. 4e). These differences might reflect MB subgroups specific tropisms. So, despite the fact that they are initially grafted at the same site, tumor masses from the various MB subgroups settle in specific locations. Interestingly, these tumor locations correlate with the cartography of MB tumors in patients established from MRI images: MB SHH are predominantly observed in the cerebellar hemispheres and MB G3/G4 at the border of the 4^th^ ventricle^3,4,25^(Fig. 4f). Moreover, these topographies are also consistent with the localizations and the early migration pathways of the putative progenitors of origin: GCP at the origin of MB SHH colonize the EGL, while early RL progenitors at the origin of MB G3 are stacked in the germinal zone which later invaginates along the 4^th^ ventricle. Alternatively, progenitors of CN and UBC, lineages of origin of more committed MBG3/G4 will migrate into the deep medio-ventral part of the primitive cerebellum and through the white matter lining the 4^th^ ventricle (Supp. Fig. 1a). Therefore, our grafting paradigm enables us to recapitulate key aspects of MB disease that relate to the site of emergence and the migratory trajectories of their embryonic cells of origin.

**Figure 4.**
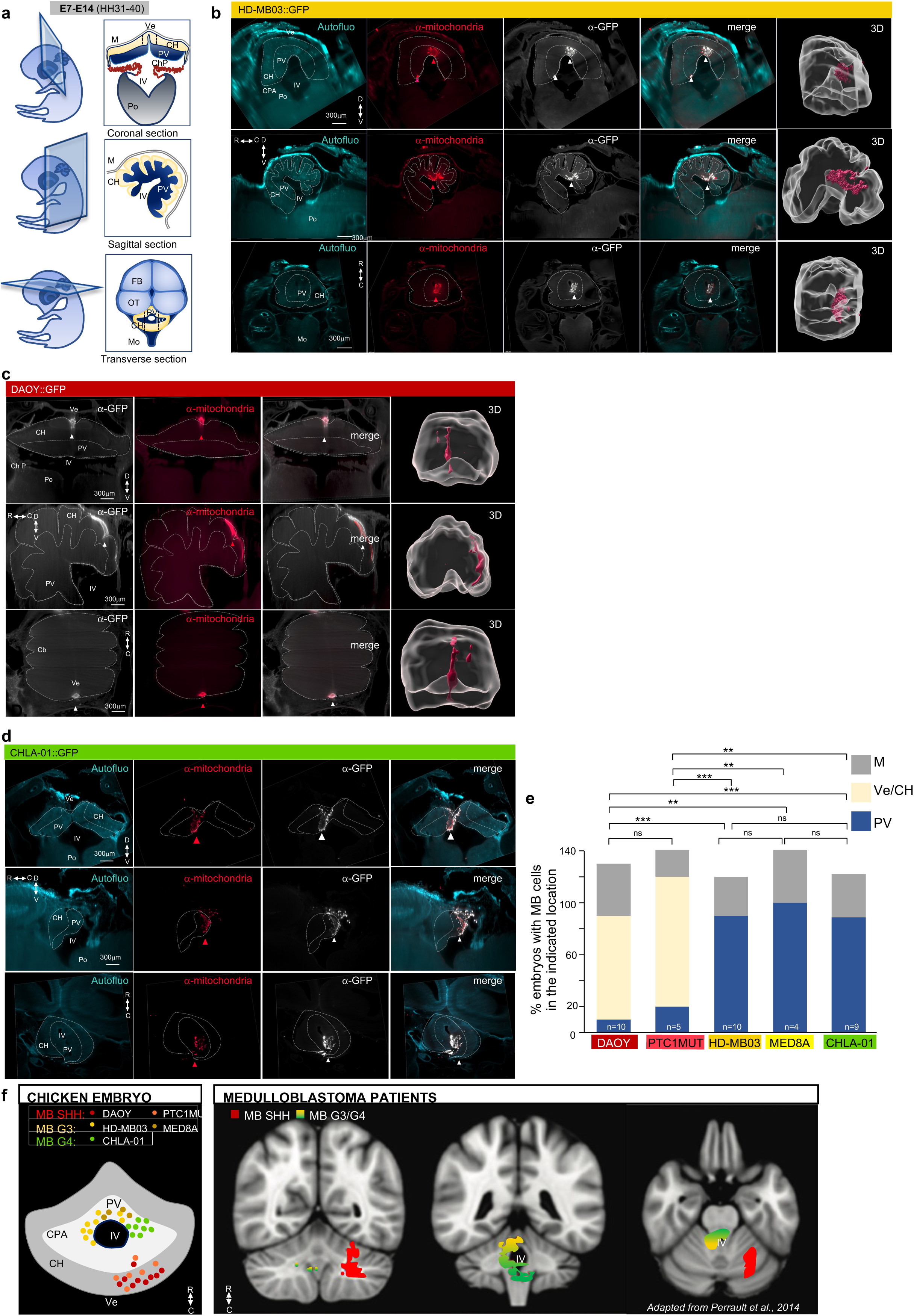
The topography of the tumors formed by the different MB subgroups reflects those observed in patients. **a.** Schematic representation of coronal (upper panel), sagittal (middle panel) and transverse section (lower panel) of the developing cerebellum of a chick embryo at late stage (E7-E14, HH31-40). **b.** Representative coronal (upper panel), sagittal (middle panel) and transverse (lower panel) optic sections of a chick embryo 10 days after the graft of HD-MB03::GFP. The embryo was in toto stained with an anti-mitochondria antibody targeting human cells and an anti-GFP antibody. Autofluorescence (autofluo) allows the visualization of structures. The whole embryo was imaged with SPIM. Arrowheads point to the tumoral masses in the paraventricular zone delimited by white dotted lines. The pial surface of the cerebellum is also delineated with white dotted lines. 3D reconstruction was performed with Imaris software to better visualize the topography of the tumor (right panel). The cerebellum outlines are represented in white and the tumor mass surface reconstruction is in pink. See also Movie S3. **c.** Representative optic sections, as in b, of a chick embryo 10 days after the graft of DAOY::GFP cells. **d.** Representative optic sections, as in b, of a chick embryo 6 days after the graft of CHLA-01::GFP cells. **e.** Quantification of the percentage of embryos at late stages (E7-E14, HH31-40) with tumor masses in the indicated location. n is the number of embryos grafted with the indicated cell lines that were analyzed. The Chi-square test was performed to compare two cell-lines and Vermis / Cerebellar Hemispheres (Ve/CH) versus Para-Ventricular (PV) location. There was no significant difference for the presence in Meninges (M). **f.** Cartography of the tumors observed in the chick embryo for each indicated cell line summarizing the results from e. Each dot corresponds to a tumor mass observed in a grafted embryo (left panel). The right panel is adapted from Perrault *et al.* 2014: the frequency of MB SHH occurrence versus combined MB G3 and G4 and the area of significant differential involvement was reported on a representative MRI optic section of a human cerebellum. M: meninges, Ve: Vermis, CH: Cerebellar hemisphere, ChP: Choroid Plexus, PV: Para-Ventricular zone, IV: fourth ventricle, Po: Pons, FB: forebrain, OT: optic tectum, Mo: Medulla oblongata, D: dorsal, V: ventral, R: rostral, C: caudal, A: anterior, P: posterior. ns : non-significant, ** p≤0.01, *** p≤0.001.

### MB cells have kept features from their lineage of origin, that they reactivate once grafted in their territory of origin in the chick embryo

Next, we further studied tumoral cells patterns in grafted embryos by confocal microscopy performed on embryo immunostained cryosections. First, we analyzed MB tumor cohesion profiles at the cellular level, which were hard to access with the eCbOTC model. Using 3D reconstruction software, we calculated the distance between each cell and its 9 nearest neighbors (d9nn) within the mass. We classified those whose distance to d9nn was 3 times superior to the average cell diameter as “isolated” cells. DAOY cells tended to form cohesive and well-delimited masses, conversely to HD-MB03 and CHLA-01 cells which showed dispersed patterns with insolated cells at the periphery of the forming tumor (Fig. 5a). These subgroup specific features might be associated with a more or less exploratory phenotype of the tumoral cells. Indeed, although grafted at the same location, HD-MB03 and CHLA-01 cells navigate toward the paraventricular zone while DAOY tumors settle at the periphery of the cerebellum hemispheres. This suggests that they adopt different navigation modes. A good indication on the migration modalities adopted by cells relies on its morphology and polarity. We thus studied the morphology of MB cells in the grafted embryos and started with HD-MB03 cells. At t=0, just after the graft in the RL vicinity, HD-MB03 cells displayed a rounded shape, as seen in culture. 24h, and particularly 48h after the graft, they had elongated, adopting a fusiform shape with extending processes (Fig. 5b). We used the method developed for the eCbOTC model (Fig. 2f) to classify morphologies in a systematic analysis of all the cells. Consistent with our previous observations, we found that HD-MB03 cells were quite rounded, displaying small processes with occasionally larger terminal “growth cone-like” structures (Fig. 5c). DAOY cells adopted a very elongated, bipolar morphology forming streams at the periphery of the mass (Fig. 5d) and CHLA-01 cells exhibited unipolar brush cell-like morphology (Fig. 5e). After quantification, we observed a significant association between DAOY cells and a bipolar morphology; and between CHLA-01 cells and a brush morphology. More generally, quantification of the total proportion of cells with specific morphologies show that CHLA-01 are prone to adopt these morphologies generally associated with differentiation features compared to HD-MB03 cells that mostly remain round like immature progenitors (Fig. 5f and Supp. Fig. 5a). Thus, morphologies of MB cells representative of the different sub-groups reveal that they might adopt a more or less differentiated state and navigation modes in the developmental sequence that is tied to their lineage of origin.

**Figure 5.**
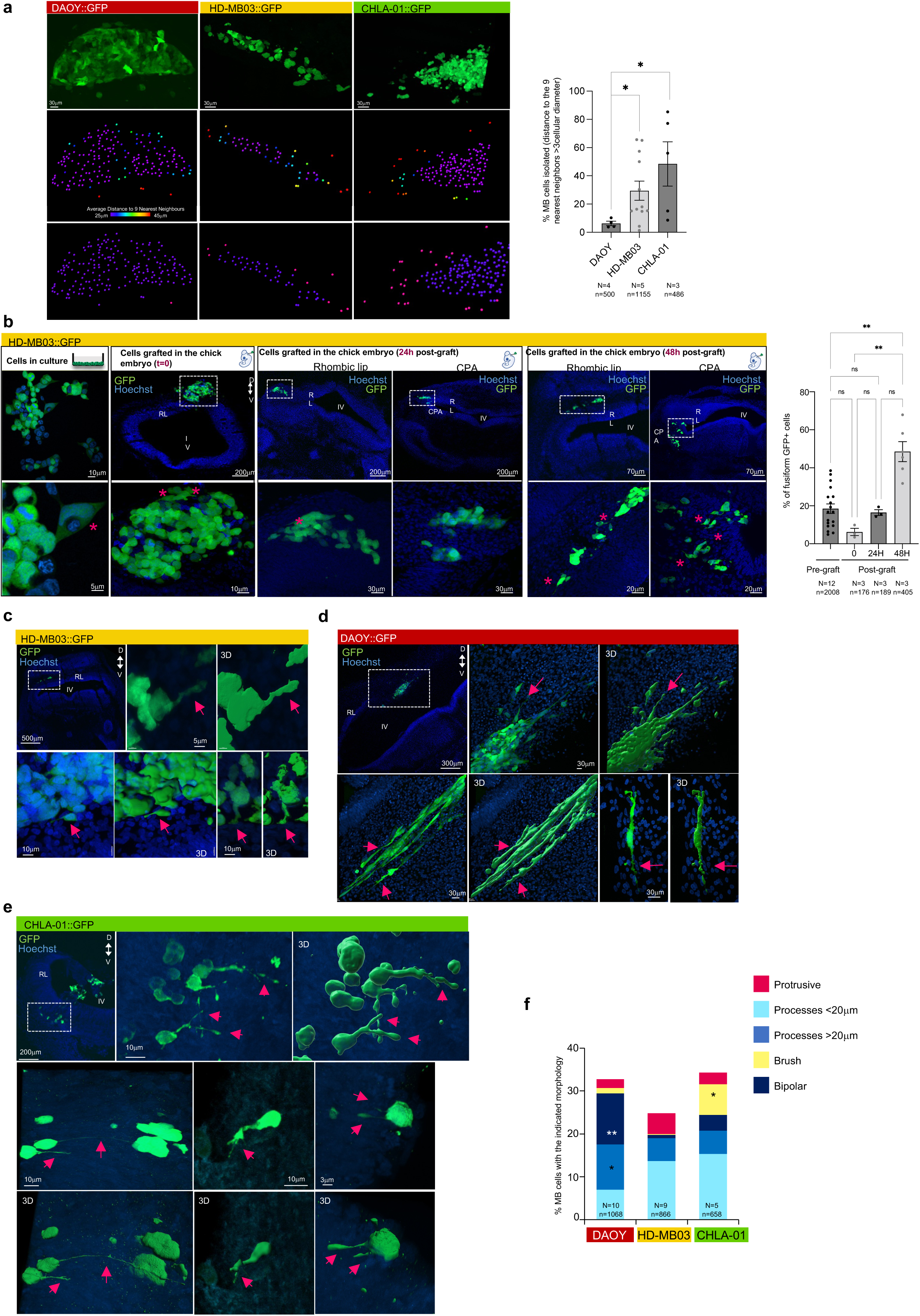
At the microscopic level, the behavior of grafted MB cells in the primitive cerebellum reflects that of their lineage of origin. **a.** Representative cryosections of embryos grafted with the indicated MB cell lines. Endogenous GFP is observed (upper panel). The cohesion of the tumor mass was evaluated by cellular segmentation and measurement of the distance of each cell to its 9 nearest neighbors (ranging from 25μm to 45μm and displayed by a spectrum look-up table (LUT)) (middle panel panel). This Imaris measurement highlights in red peripheral cells isolated from its 9 nearest neighbors from a distance > 3ad (ad: average diameter of the cell) in pink (lower panel). Quantification is shown on the right. The number of embryos (N) and cells (n) analyzed are indicated. A Mann-Whitney test was performed comparing each HD-MB03 and CHLA-01 to DAOY condition. **b.** Representative coronal cryosections of embryos grafted just after the graft (t=0), 24h and 48h of HD-MB03::GFP. Endogenous GFP of grafted cells and Hoechst staining of nuclei are shown. Tumors in the RL (RL) and in the (CPA) are observed. The lower panels are enlargements of the upper panels. Cells cultured on slices were observed and the number of fusiform cells were counted (pre-graft, N=12 different cells batches, n=2008 counted cells). GFP positive cells were counted on cryosections of grafted embryo at the indicated time post-graft (N: number of embryos, n: number of counted cells). Similarly, the number of fusiform cells was counted for each condition (representative fusiform cells are labeled with a pink asterisk). A Kruskal-Wallis test was performed. Error bars indicate SEM. ns: non-significant, * p≤0.05, ** p≤0.01, *** p≤0.001, **** p≤0.0001. **c.** Representative coronal cryosections of embryos grafted with HD-MB03::GFP as in b. In the upper left panel, a representative image of the primary tumor in the vicinity of the RL is shown in the white dotted line square. Higher resolution acquisitions are displayed in the adjacent panels. Extended processes are highlighted by pink arrows. Imaris surface reconstruction (3D) allows a better visualization of the processes. **d.** Representative coronal cryosections of embryos grafted with DAOY::GFP as in c. Characteristic bipolar shaped cells are highlighted by pink arrows. **e.** Representative coronal cryosections of embryos grafted with CHLA-01::GFP as in c. Characteristic bi-ramified, brush cells and long processes are highlighted by pink arrows. **f.** Striking morphological features were quantified for the different MB subgroups cell lines. They were calculated as a percentage of the total number of cells analyzed. The number of embryos (N) and cells (n) analyzed are indicated. For each morphology, a Mann-Whitney test was performed to compare two cell lines at a time. Only statistically significant features specific to one condition are indicated. * p≤0.05, * p≤0.05, ** p≤0.01.

### The impact of the embryonic microenvironment on the transcriptome of MB cells

In order to characterize the molecular dialogues between tumor cells and their microenvironment, we performed a bulk RNA Seq analysis. We focused on the HD-MB03 cell line and compared the transcriptomic content of pre-grafted cells and of cells that experienced the miroenvironment provided by the cerebellar primordium of chick embryos during 3 days. We grafted 3 different batches of “pre-graft” cells at E3 and microdissected the corresponding 3 batches of embryos at E6 to extract the tumor masses formed in the vicinity of the RL (“post-graft” triplicates). The tumor masses were dissociated, the GFP-positive cells were sorted and sequenced (Fig. 6a). We confirmed the absence of batch effect in the analysis by Principal Component Analysis (PCA) (Supp. Fig. 6a).

**Figure 6.**
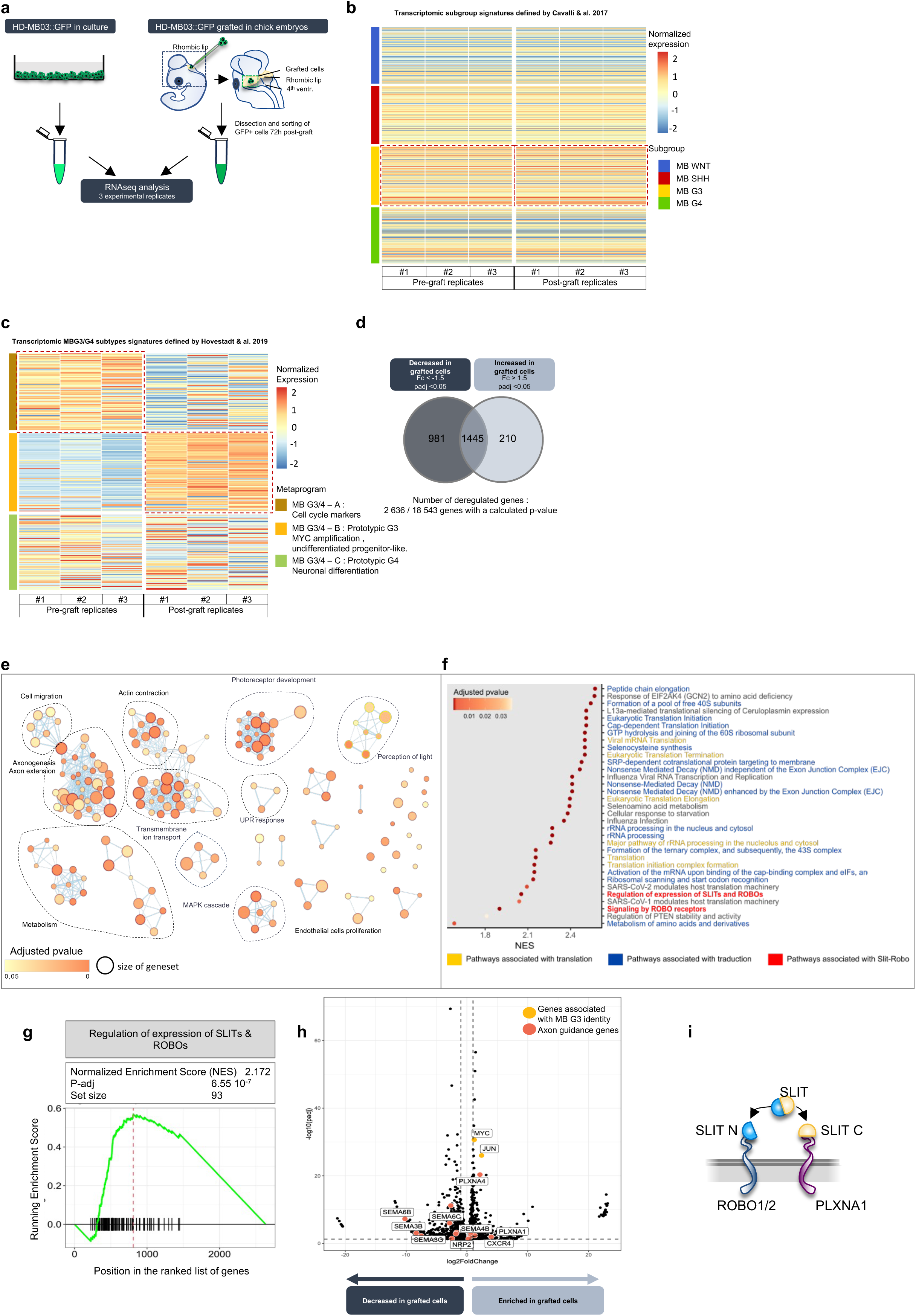
Upon the graft in the chick primitive cerebellum, MB G3 activate transcriptomic programs from MB patients and the canonical SLIT signaling pathway. **a.** Overview of RNA seq analysis comparing HD-MB03::GFP naïve cells (before the graft) versus microdissected tumor masses 72h post-graft. Both experimental conditions were sequenced in triplicate. **b.** Heatmap of mRNA expression for upregulated and downregulated MB subgroups classifying gene sets published by Cavalli *et al.,* 2017, in HD-MB03::GFP cells before and after the graft. The z-score for each transcript is color-coded. Dotted square lines highlight the most significantly enriched signature in each condition. Both experimental conditions were sequenced in triplicate. **c.** Heatmap of mRNA expression for upregulated and downregulated MB G3/4 metaprograms published by Hovestadt *et al.,* 2019, in HD-MB03::GFP cells before and after the graft. Dotted lines highlight the most significantly enriched signature in each condition. The z-score for each transcript is color-coded. **d.** Venn diagram indicating the number of significantly differentially expressed transcripts compared to the control condition of pre-grafted cells. **e.** Over-representation enrichment map based on significantly deregulated genes (Abslog2fc>1.5, padj≤0.05) from the Gene Ontology Biological Processes database. Nodes (circles) represent enriched pathways, their size corresponds to the size of the gene set and the line width to the number of genes that two pathways have in common. The overlap cutoff is 50%. **f.** Gene set enrichment analysis (Reactome database, ClusterProfiler, padj≤0.05) in post-versus pre-graft HD-MB03. **g.** Enrichment plots used to calculate the normalized enrichment score (NES) of “Regulation of expression of SLITs and ROBOs”) gene set (padj and set size are indicated). **h.** Volcano plot highlighting significantly decreased or enriched genes. Vertical dotted lines represent (Abslog2fc=1.5) and horizontal ones (padj≤0.05). Genes characteristic of MB G3 and axon guidance genes (see Supp. Table 2) are highlighted. **i.** Schematic representation of SLIT cleavage and binding of its N-terminal part (SLIT N in blue) with ROBO1/2 and its C-terminal fragment (SLIT C in yellow) to PLXNA1.

First, we studied whether the embryonic microenvironment modifies MB subgroup identity. With this aim, we extracted consensual subgroup transcriptomic signatures generated from 763 MB patient samples from the 4 MB subgroups by F. Cavalli and colleagues (GSE85217)^26^. By analyzing these signatures in our dataset, we found that both pre- and post-graft replicates best matched with MB G3 signature, which confirms that the cerebellar embryonic microenvironment maintains MB cell identity (Fig. 6b). We then extracted signatures revealed in MB patients G3 and G4 from another cohort^27^. We observed that, within the cerebellum primordium, MB cells switched from a proliferative cell cycling toward an undifferentiated progenitor-like transcriptional state, prototypical of MB G3 associated with MYC amplification (Fig. 6c). Interestingly, the cerebellum microenvironment thus initiated transcriptional programs known to be active in MB patients.

Second, we analyzed the genes that were differentially expressed between pre-and post-grafted cells and found 2636 out of 18453 genes having significant differences (padj<0.05). 1191 of them displayed strong expression levels changes (absval(LOG2 Fc) >1.5), of which 981 had lower expression and 210 that were up-regulated after the graft (Fig. 6d). Among the most significantly upregulated genes, we found MYC^28^, JUN^29^ and CXCR4^30^ already characterized as markers of aggressive MB G3 (Supplementary Table 1). Then, to study the biological processes regulated by the embryonic cerebellum microenvironment, we performed an over-representation analysis of the 1191 genes using Gene Ontology Biological Process database. Significantly over-represented pathways were related to MB G3 characteristic processes such as general metabolism, the MAPK cascade, and pathways related to the development of photoreceptors^31^ which is also linked to the MB G3 identity (Fig. 6e). Others were related to neuronal identity (axonogenesis or transmembrane ion transport) and also to cellular migration, consistent with the observed behaviors and morphologies of grafted MB cells (Fig. 5bcf). Next, to refine this pathway analysis, we performed Gene Set Enrichment Analysis on the total of 2636 significantly deregulated genes using Reactome database. Within the significantly enriched pathways (padj<0.05) most of them were associated with translation, which is typical of MB G3 (Fig. 6f). Notably, two signatures related to SLIT-ROBO canonical axon guidance pathway appeared among the most significantly enriched pathways. Both had positive Normalized Enrichment Score (NES, 2.172 and 2.099, Fig. 6g and Supp. Fig. 6b) suggesting a significant deregulation of the signature genes in grafted cells. To closely assess the transcriptional regulations reflecting the morphological features and navigation modes adopted by grafted cells, we studied the dynamic of expression of a homemade list of axon guidance related genes (Supplementary Table 2). We found several guidance receptors and ligands whose expression was modified in grafted cells, thus reflecting that MB cells use guidance cues present in the embryonic environment, SLIT being among them (Fig. 6h).

SLIT is a secreted protein of 200kDa. Its cleavage generates a large N-terminal (SLIT N) fragment of 150kDa and a smaller C-terminal one of 50kDa (SLIT C) (Fig. 6i). SLIT N binds to ROBO1/2 receptors and its role as a major repulsive cue has been revealed by studies of neurons axonal navigation of neurons (for review^32^). SLIT C has long been considered as inactive until it was demonstrated to induce a repulsive signal via PLXNA1 receptor^33^. Since its discovery as an axon guidance signal in Drosophila, SLIT signaling has been found to be involved in many processes linked with cytoskeleton remodeling, cellular migration, organogenesis, angiogenesis and maintenance of stemness during embryonic physiological development but also in pathological contexts and particularly in various cancers (for review^34,35^). Therefore, we hypothesized that the observed transcriptional regulations of SLIT signaling in grafted cells could mediate a range of behaviors during the early steps of MB tumorigenesis.

### SLIT2 is expressed in the RL during early cerebellum development and MB G3 cells are able to perceive it, via ROBO1 and PLXNA1

A previous pioneering study showed that SLIT2 is expressed at the edge of the RL during chick cerebellum development and suggested that it might repel the progenitors expressing ROBO1/2 out of the germinal zone^8,36^. In situ hybridization performed on rat embryos also revealed the expression of SLIT2 in the Purkinje cell layer^37^. To assess if these expression profiles are conserved in the chick embryo at early stages and if SLIT2 could be exploited by grafted MB cells, we achieved Hybridization Chain Reaction (HCR) RNA Fluorescence In Situ Hybridization (FISH) combined with clearing and SPIM imaging of the embryo^38^. We first set up the staining with a *Pax6* probe to observe the migration of RL-derived progenitors^39^. At E3, the Pax6 probe stained the progenitors that started to emerge from the germinal zone at the edge of the RL to undergo a rostral subpial migration (Fig. 7a). While we found the highest *Slit2* expression in the spinal cord, *Slit2* RNA was additionally detected at the edge of the RL (Fig. 7b). At E4, *Slit2* expression was also visible in the roof of the 4^th^ ventricle from which the choroid plexus would emerge (Fig. 7c). At E6, due to RL growth, we proceeded to sagittal cryosections stained with HCR-RNA FISH and observed that *Pax6* expressing progenitors still emerged from the RL (Fig. 7d) and that *Slit2* was still expressed in the germinal zone and in the choroid plexus anlage (Fig. 7e). We also observed its expression in the Purkinje cell layer (Fig. 7f). Higher resolution confocal imaging revealed that *Slit2* seemed to be expressed along the immature Bergman glial cells fibers^40^. Putative exposition of SLIT2 by glial processes has already been reported in the floor plate and demonstrated to drive and channel the navigation of ROBO1 and PLXNA1 expressing axons via repulsion ^41^. Overall, we observed high *Slit2* expression at the edge of the RL lining ventrally the migration of Pax6 positive progenitors. The complementary patterns of expression of *Pax6* and *Slit2* suggested that *Slit2* could repel the progenitors out of the germinal zone, constraining and channeling their migration along the pial surface (Fig. 7g). Finally, we observed that *Slit2* expression was maintained at the edge of the RL until the late progenitors emerge at E8 (Fig. 7h). We also analyzed the single-cell RNAseq data set from Aldinger and colleagues (2021) encompassing 9-20 PCW human cerebellar cells^42^ (Fig. 7i). We subsetted RL-derived neuronal progenitors (Fig. 7j) and observed that SLIT2 expression was high in human RL early progenitors (Fig. 7k) as demonstrated in the chick embryo by our FISH staining. In comparison, SLIT1 and SLIT3 were much less expressed in the human early cerebellar progenitors suggesting that SLIT2 is probably the main ligand of the pathway functionally involved in this context (Supp. Fig. 7a).

**Figure 7.**
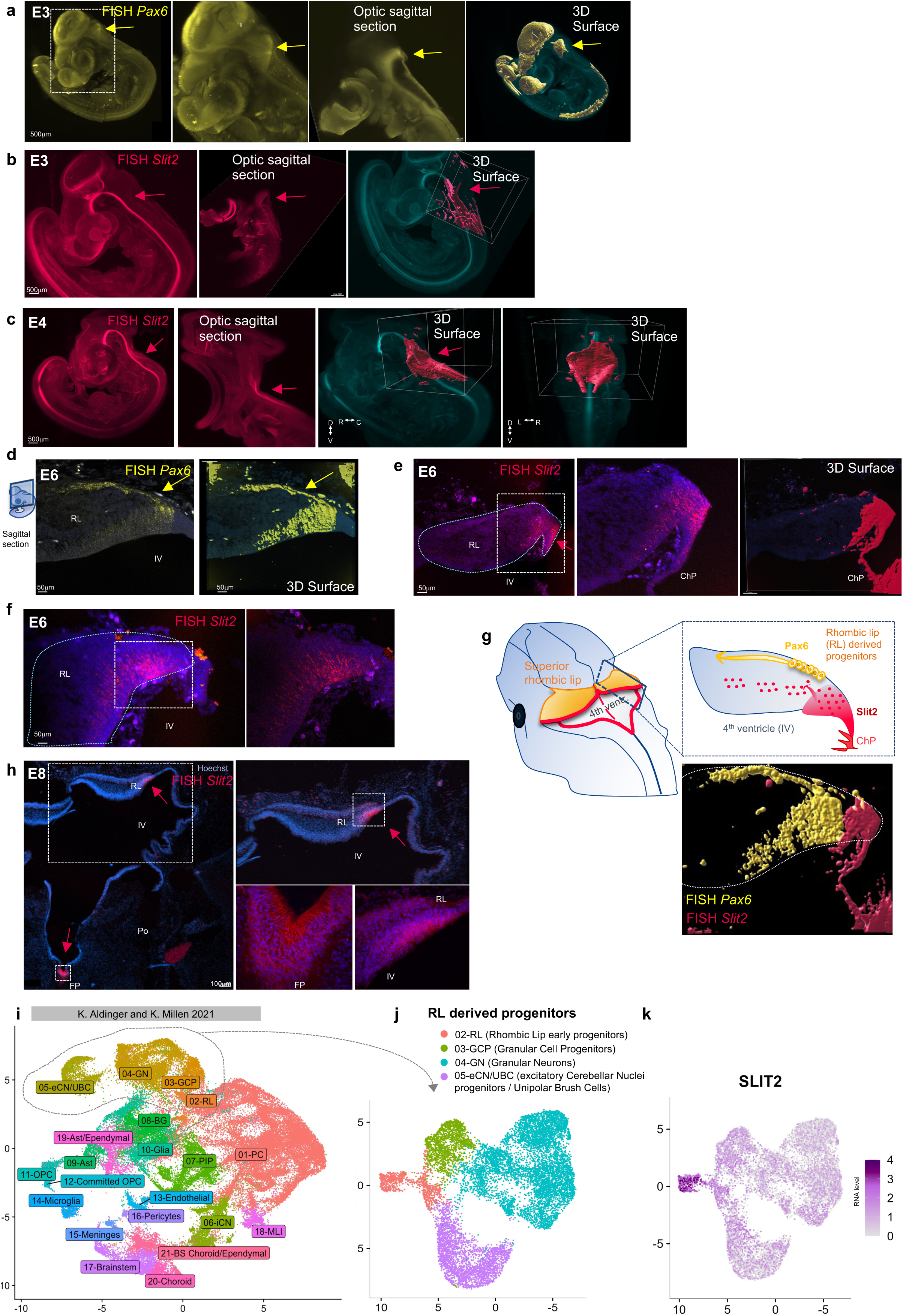
SLIT2 is expressed in the primitive RL when progenitors emerge from the germinal zone. **abc.** In toto Hybridization Chain Reaction (HCR) RNA Fluorescence In Situ Hybridization (FISH) combined with clearing and SPIM imaging of E3 (a and b) and E4 (c) chick embryos with *Pax6* (a) and *Slit2* (b and c) probes. PAX6 is a marker of RL-derived progenitors. Staining in the RL is indicated by arrows. 3D surface reconstruction with Imaris software allows a better visualization of the staining in the RL (a, b, c) and in the roof of the 4^th^ ventricle (c). **def.** HCR-FISH performed on cryosections of E6 embryo with *Pax6* (d) and *Slit2* (ef) probes. The RL is delimited by blue dotted lines. The square delimited area (white dotted lines) in the left panel is enlarged on the right. **g.** Schematic representation of the early developing embryo. The RL is labeled with orange and a sagittal section is enlarged displaying *Pax6* positive progenitors (in yellow) emerging from the *Slit2* positive germinal zone (in pink). The ventral expression of *Slit2* is indicated by pink dots. In the lower panel *Pax6* and *Slit2* staining surface reconstruction at E6 are merged and show a striking complementarity of expression. **h.** HCR-FISH performed on cryosections of E8 embryo with *Slit2* probe. An enlargement of the square delimited areas is shown for both the RL and the FP. **ijk.** Analysis of the expression of SLIT2 in single cell RNASeq data of the human developing cerebellum (Aldinger et al., 2021). UMAP visualization of 67174 human cerebellar nuclei colored by cluster identity from Louvain clustering and annotated on the basis of marker genes (9-20 PCW) (i). RL-derived progenitors were subset in a new UMAP (j). The average expression of SLIT2 in this UMAP embedding is shown in (k). The color indicates log-normalized gene expression. RL: RL, IV: 4^th^ ventricle, ChP: Choroid Plexus, FP: Floor Plate, Po: pons, GCP: Granular Cell Neurons, GN: Granular Neurons, eCN/UBC: excitatory Cerebellar Nuclei/Unipolar Brush Cells. eCN (notation from Aldinger et al,) correspond to CN in the text.

Next, we studied if MB cells might take advantage of this SLIT2 RL source. We first analyzed our in-house collection of 341 MB patient samples (Bernardi *et al.,* under revision) and 16 MB cell lines and combined transcriptomic (RNA-seq), methylomic (850K), proteomic (MS) and phospho-proteomic (MS). We observed that ROBO1 and PLXNA1 receptors were expressed in MB G3 patients both at the RNA (Fig. 8ab) and protein level (Fig. 8cd) and in all MB G3 subtypes (Supp. Fig. 7bcde). We confirmed this expression at the RNA level in transcriptomic data set from a public cohort of 763 MB patients’ tumors provided by Cavalli and colleagues^26^ (GSE85217, Supp. Fig. 7fg). So, we speculated that MB G3 cells might be able to perceive SLIT ligands. We consistently found, both in our in-house cell lines collection (Fig. 8efgh) and in public transcriptomic data sets (Supp. Fig. 7hi), that parental MB G3 HD-MB03 cells displayed high ROBO1 and PLXNA1 expression. Particularly, ROBO1 is highly expressed in HD-MB03 cells compared to MB G4 CHLA-01. We also carried out RT-Q-PCR and immunostaining to ensure that these expressions were still detected in our GFP-positive HD-MB03::GFP cells. We found that both receptors were expressed at high mRNA levels, compared to control neuroblastoma SHEP cells and to CHLA-01::GFP cells which express a high level of ROBO2 (Fig. 8i). ROBO1 and PLXNA1 proteins are specifically expressed at the cell membrane of HD-MB03, as expected for receptors of extracellular ligands ^43,44^ (Fig. 8j).

**Figure 8.**
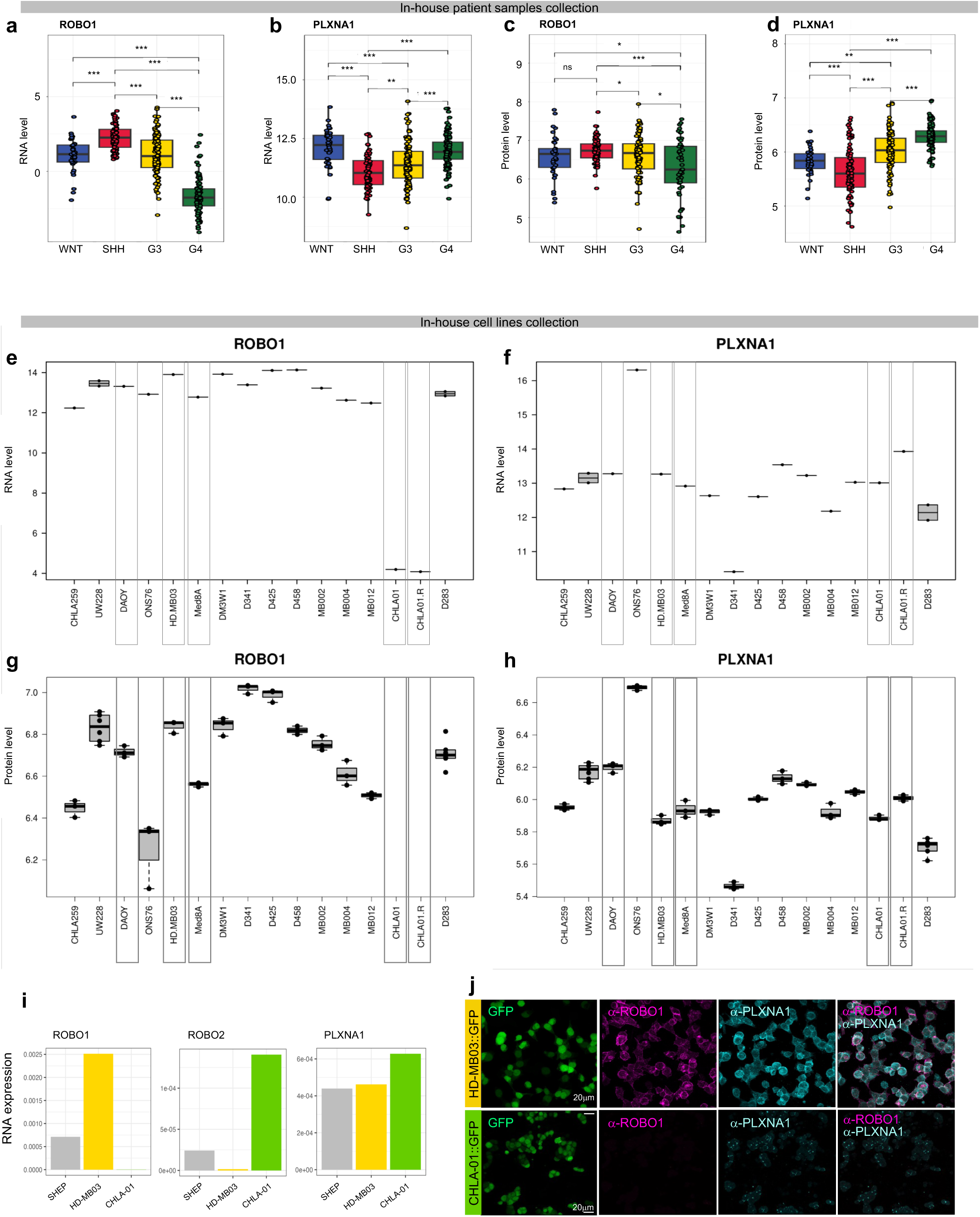
SLIT receptors ROBO1 and PLXNA1 are expressed in MB G3 tumors and cell lines. **abcd.** Average expression, at the mRNA (ab) and protein level (cd) of ROBO1 (ac) and PLXNA1 (cd) in MB tumors from our in-house cohort. **efgh.** Average expression, at the mRNA (ef) and protein level (gh) of ROBO1 (eg) and PLXNA1 (fh) in MB cell lines from our in-house collection. **i.** RT-Q-PCR measure of ROBO1, ROBO2 and PLXNA1 mRNA in Neuroblastoma SHEP control cells, CHLA-01::GFP (MB G4) and HDMB-03::GFP (MB G3 cells). The ratio between mRNA level and the housekeeping gene GAPDH mRNA level, i.e., the 2^ΔCt^ raw value = 2^-(Ct Gene of Interest – Ct GAPDH)^, is color coded as indicated. **j.** Immunostaining performed on HD-MB03::GFP and CHLA-01::GFP cells grown on slices with α-ROBO1 and α-PLXNA1 antibodies.

### SLIT signaling loosens cell-cell cohesion and might confer infiltrative properties to MB G3

Next, we investigated whether SLIT signaling modulates the properties of MB G3 cells, considering its pleiotropic functions^32^. We first tested the impact of SLIT signaling on spheroids of HD-MB03 cells in a hanging-drop cohesion assay. We observed that addition of both SLIT2 N and SLIT2 C recombinant fragments to the medium induced the cohesion release of the cells within the aggregates (Fig. 9a). In contrast, SLIT2 fragments had no impact on tumor cells survival (Supp. Fig. 8a) or proliferation (Supp. Fig. 8b). We then invalidated both ROBO1 and PLXNA1 by siRNA in HD-MB03 cells (Supp. Fig. 8c) and were able to reverse the cohesion release effect of SLIT2 fragments on MB cells (Fig. 9b).

**Figure 9.**
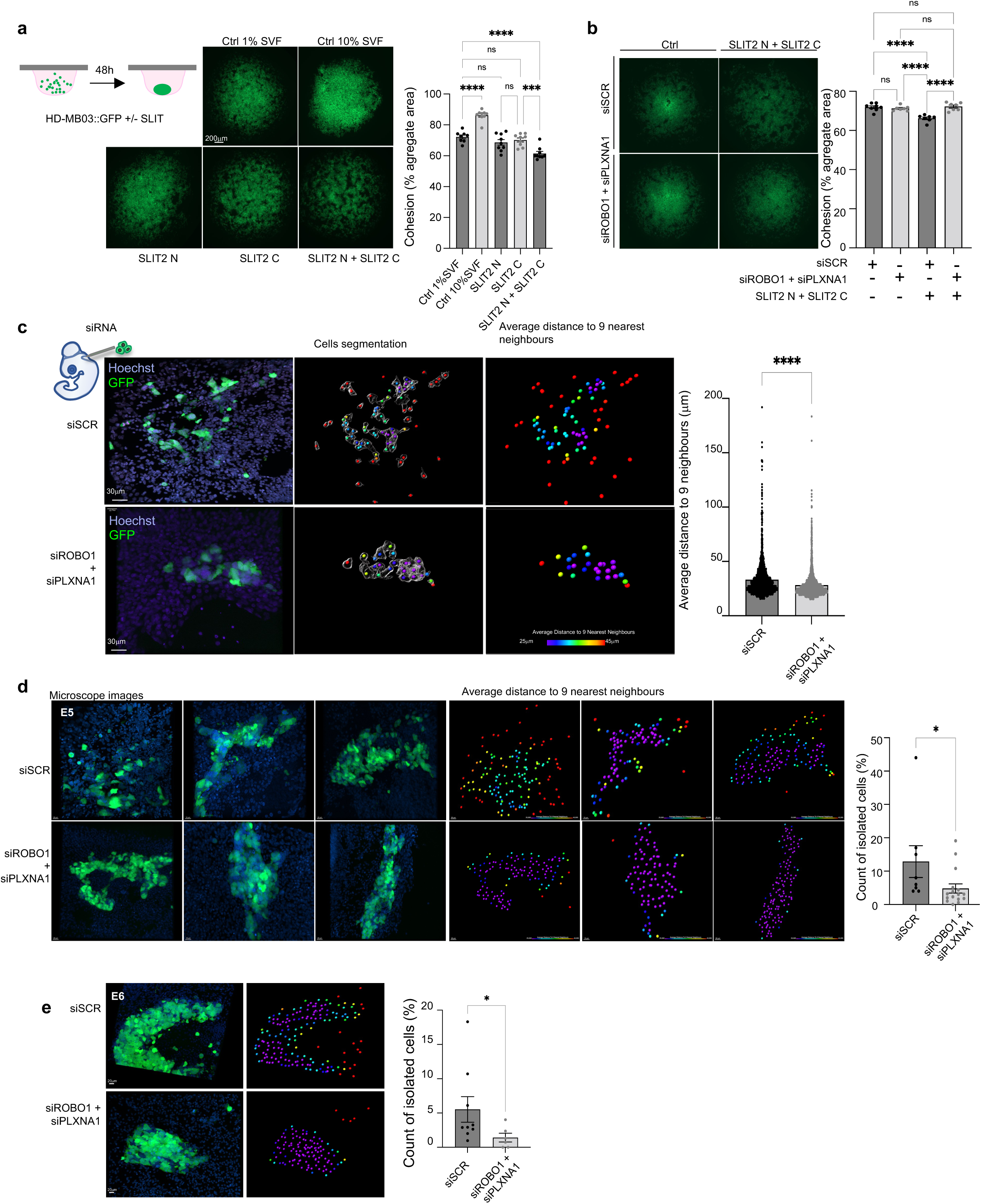
SLIT loosens cell cohesion and favors the exploratory behavior of MB G3 cells via ROBO1 and PLXNA1. **ab.** Hanging-drop assays performed on HD-MB03::GFP to monitor the formation of aggregates and their cohesion. Cellular suspensions were cultured with or without SLIT recombinant fragments SLIT N and SLIT C (2mg/ml final concentration each) resuspended in RPMI medium with 1% SVF. RPMI with 10% SVF was used as an aggregation positive control (a). The cohesion release induced by SLIT fragments was rescued by invalidation of ROBO1 and PLXNA1 by siRNA (siROBO1 + siPLXNA1) (b). Representative images of the aggregates are shown on the left. The analysis was performed at least 3 times with at least 7 aggregates per condition. The Anova test was applied. **cd.** Representative cryosections of embryos grafted with HD-MB03::GFP transfected with siScramble (siSCR) or siRNA targeting ROBO1 and PLXNA1 (siROBO1 + siPLXNA1), 48h post-graft (E5). Endogenous GFP is observed (left panel). For c and d, different representative sections are shown for each condition. The cohesion of the tumor mass was evaluated by cellular segmentation (c, middle panel) and measurement of the distance of each cell to its 9 nearest neighbors (d9nn) (ranging from 25mm to 45mm and displayed by a spectrum look-up table (LUT)) (cd, right panel). The average d9nn distances of each considered cells were cumulated on a diagram (c). More precisely, this measurement performed by Imaris software highlights in red the peripheral cells isolated from its 9 nearest neighbors from a distance > 40mm (corresponding to 3x average diameter of HD-MB03 cells). These isolated cells were counted and their percentage over the total number of cells counted was calculated (d). For c and d, different grafted embryos (N) with different cryosections (cs) and a different number of cells (n) were considered. siSCR: N=4, cs=9, n=1901, siROBO1 + siPLXNA1 : N=4, cs=6, n=999. **e.** As in d, isolated cells were counted in chick embryos grafted with siSCR or siROBO1 + siPLXNA1, 3 days post-graft (E6). SiSCR: N=3, cs=8, n=889, siROBO1 + siPLXNA1: N=6, cs=16, n=2142. The Mann-Whitney test allowed the comparison of both conditions. Error bars indicate SEM. ns: non-significant, * p≤0.05, **** p≤0.0001.

We then studied the role of SLIT signaling, in vivo, using our chick embryo model, by grafting HD-MB03 cells, invalidated or not for SLIT receptors, in the chick embryo. Two days after the graft, at E5, we collected the embryos and carried out serial cryosections. We selected the cryosections encompassing the GFP-positive grafted cells forming masses in the vicinity of the RL. Whereas HD-MB03 cells transfected with control siSCR formed poorly cohesive and dispersed tumor masses, as already observed with non-transfected cells (Fig. 5a), we observed that HD-MB03 cells invalidated for SLIT receptors (siROBO1 + siPLXNA1) formed more cohesive masses (Fig. 9c). In order to quantify these observations, we proceeded to cell segmentation and measured the distance of each cell to its 9 nearest neighbors (d9nn) with Imaris software. As in Fig. 5a, this measure was used as a read out of tumor cells cohesion. We observed a significant difference of average d9nn between the masses formed by HD-MB03 cells transfected with siSCR control (average d9nn = 33,2μm) and those transfected with siRNA invalidating both receptors (siROBO1 + siPLXNA1, average d9nn = 28,2 μm) (Fig. 9c). This result supports that SLIT signaling confers less cohesive and more motile behaviors to HDMB-03. The d9nn quantitative method also allowed us to analyze isolated and dispersed cells (d9nn > 40μm). Strikingly, these peripherally isolated cells were significantly more numerous in the control tumor masses while they were absent or rare in the tumor masses deprived of SLIT signaling (Fig. 9d). We verified that this observation did not reflect a delay in the formation of the tumors and carried out a similar analysis one day later (E6) when the tumors were bigger. At this stage also, we observed more isolated peripheral cells in the condition of active SLIT signaling (Fig. 9e).

### SLIT signaling is associated with a poor prognosis in MB G3 patients

These SLIT-mediated putative spreading properties could contribute to the aggressivity of MB cells. We thus examined whether receptor expression levels in MB tumors discriminate the outcome of patients in clinic. We segregated ROBO1^high^PLXNA1^high^ versus ROBO1^low^PLXNA1^low^ tumors among the 87 MB G3 of our in-house collection of MB tumors. We observed that ROBO1^high^PLXNA1^high^ had a tendency to be associated with bad prognosis especially at the transcriptomic level, but the result was not significant (Fig. 10a). So, we turned to the publicly available survival data obtained from a collection of 763 samples (F. Cavalli – GSE85217)^26^. We segregated ROBO1^high^ PLXNA1^high^ versus ROBO1^low^ PLXNA1^low^ tumors among the 144 MB G3 and observed a significant association of high ROBO1 and PLXNA1 expression with a poor prognosis (Fig. 10b). ROBO1 or PLXNA1 expression separately was not associated with a significant worth overall survival neither in our in-house cohort (Supp. Fig. 9ab), nor in the Cavalli’s one (Supp. Fig. 9cd). Interestingly, the association of ROBO1^high^ PLXNA1^high^ expression with bad prognoses seemed to be specific of MB G3, as it was not significant for MB SHH (Supp. Fig. 9e) or MB G4 (Supp. Fig. 9f) tumor groups.

**Figure 10.**
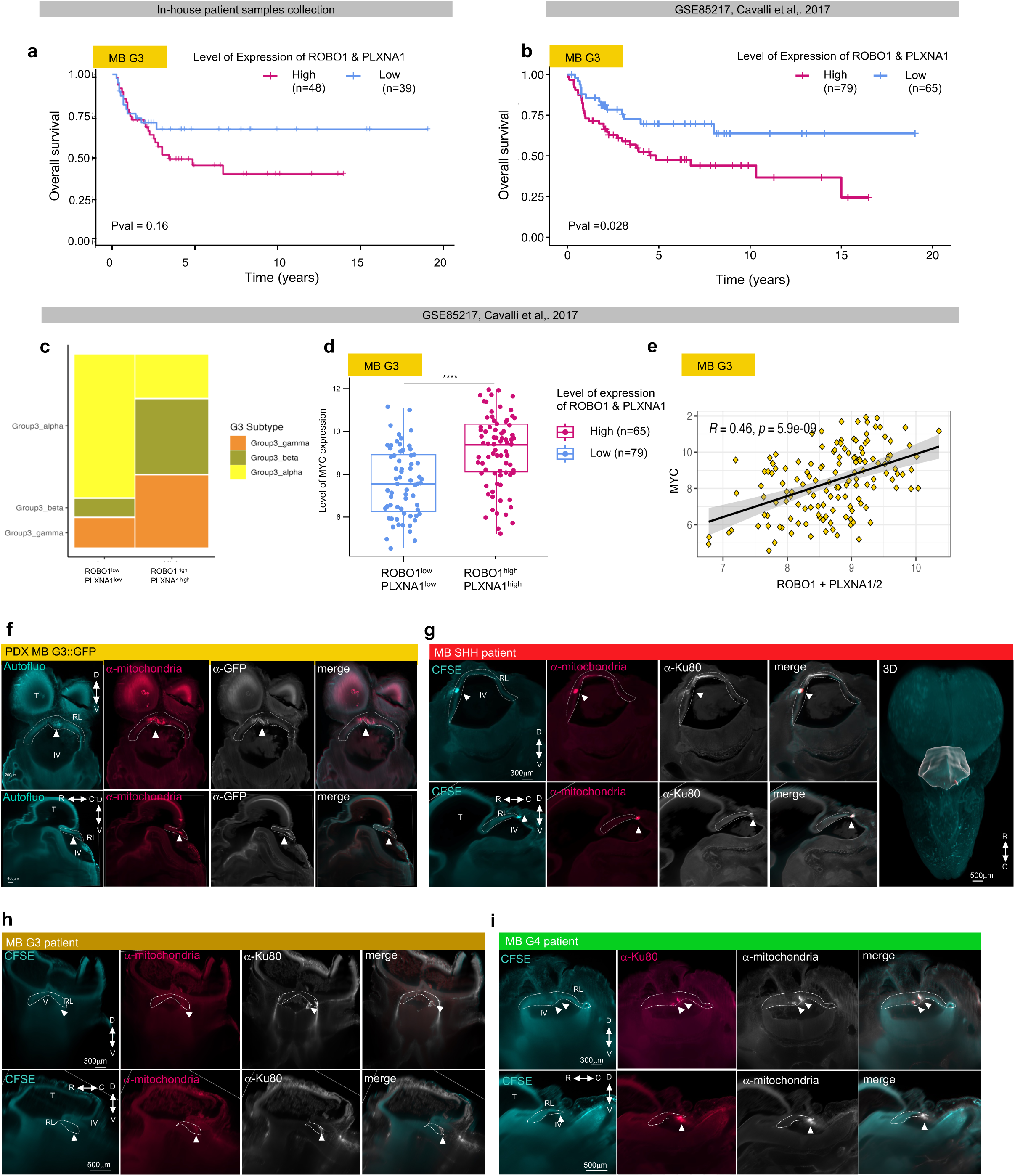
SLIT signaling is still active in human tumors and associated with aggressivity. Patient samples form tumor masses in the chick model. **a.** Kaplan-Meier overall survival curve was performed on our in-house patient samples collection, on MB G3 groups based on ROBO1 and PLXNA1 mRNA level. Mean ROBO1 and mean PLXNA1 are calculated as normalized log2 and they determine the cut-off boundary to segregate tumors in ROBO1^high^PLXNA1^high^ (High, n=48)) or ROBO1^low^PLXNA1^low^ groups (Low, n=39). The calculated p value (pval) of the Cox Proportional Hazard Model demonstrates that the two groups are not statistically different. **b.** Kaplan-Meier overall survival curve was performed on groups defined as in (a) on MB G3 tumors from the cohort of Cavalli et al., 2017, based on ROBO1 and PLXNA1 mRNA level. Mean ROBO1 and mean PLXNA1 are calculated as normalized log2 and they determine the cut-off boundary to segregate tumors in ROBO1^high^PLXNA1^high^ (High, n=79)) or ROBO1^low^PLXNA1^low^ groups (Low, n=65). The calculated p value (pval) of the Cox Proportional Hazard Model demonstrates that the two groups are statistically different. **c.** The two groups defined in b were classified based on their subtype (MB G3α (Group3_alpha), MB G3α (Group3_beta), MB G3γ (Group3_gamma)). **d.** The two groups defined in b were evaluated for their expression of MYC associated with aggressivity in MB G3. A T-test was performed between both conditions. **** p≤0.0001. **e.** Pearson correlation analysis comparing MYC and ROBO1+PLXNA1 expression. Coefficient of determination R2 and the pvalue P are indicated. **f.** Representative coronal (upper panel) and sagittal (lower panel) optic sections of a chick embryo 72h post-graft of PDX::GFP MBG3 dissociated cells. The chick embryos were stained in toto with an anti-mitochondria antibody targeting human cells and an anti-GFP antibody. Autofluorescence (autofluo) allows the visualization of structures. The whole embryo was imaged with SPIM. Arrowheads point to the tumoral masses in the RL (RL) delimited by white doted lines. α-GFP staining of both murine and human cells corresponds roughly to the α-mitochondrial staining of human cells. This observation suggests that the PDX sample is composed predominantly of human patient cells. **ghi.** Representative coronal (upper panel) and sagittal (lower panel) optic sections of a chick embryo 72h post-graft of dissociated cells extracted from MB SHH (g), MB G3 (h) and MB G4 (i) patient samples (see Table 3). Dissociated cells were stained with CFSE before the graft, although with different efficacy depending on the samples. To ensure tumor cells observation, chick embryos were stained in toto with two antibodies targeting specifically human antigens: α-mitochondria and α-Ku80. The whole embryo was imaged with SPIM. Arrowheads point to the tumoral masses in the RL (RL) delimited by white doted lines. 3D reconstruction was performed with Imaris software to better visualize the topography of the tumor. The cerebellum outlines are represented in white and the tumor mass surface reconstruction is in pink (g).

We also observed that the ROBO1^high^ PLXNA1^high^ contingent of samples were not restricted to a MB G3 subtype but rather found in G3α, and slightly more in G3β and G3γ (Fig. 9c)^26^. They were associated with a significantly higher expression of MYC, a prototypical hallmark of MB G3 aggressivity (Fig. 9de). Of note, the expression of ROBO1 and PLXNA1 alone was also slightly higher in G3β and G3γ subtypes (Supp. Fig. 10ab) and significantly associated with MYC expression, yet with lesser significance than ROBO1^high^ PLXNA1^high^ samples (Supp. Fig. 10cdef). These data can be specified in our own data set. ROBO1^high^ PLXNA1^high^ samples were associated with an equal proportion of G3α, G3β and G3γ subtypes (Supp. Fig. 10g). They were found more significantly among proteomic defined subtype G3_a (Bernardi *et al.,* under revision) (Supp. Fig. 10h) and methylome defined subtype I^45^ (Supp. Fig. 10i). The expression of ROBO1 and PLXNA1 separately did not show any obvious differences among these subtypes (Supp. Fig. 10jkl). As observed in Cavalli’s cohort, ROBO1+PLXNA1 expression was weakly yet significantly correlated with MYC, in MYC amplified and MYC non-amplified tumors (Supp. Fig 10m), while the corresponding correlations were weaker or less significant for each receptor separately (Supp. Fig 10no).

Altogether, these results suggest that SLIT signaling via both receptors ROBO1 and PLXNA1 confers more aggressive behaviors to MB G3 tumors.

### PDX and patient samples of all MB subgroups establish tumors in the avian embryo model

Finally, we investigated whether our model could be extended to patient derived samples and constitute a new preclinical model to study MB. We first studied a patient-derived xenograft (PDX) from MB G3 cells isolated from murine PDX model. In this model, GFP was expressed in tumor cells by lentiviral infection. GFP positive tumors were dissected out of the mice, dissociated and frozen in DMSO. Dissociated cells were then implanted into batches of chick embryos. Three days later, dissected embryos were subjected to in toto α-GFP and α-mitochondria staining, clearing and SPIM imaging. We observed the presence of a tumor mass – although much smaller than those obtained with cell lines - in the vicinity of the RL in most of the grafted embryos. The staining pattern of α-mitochondria antibody targeting only human cells was consistent with that of GFP labeling both human cancer cells and murine stroma, indicating that the avian tumors were not significantly contaminated by murine cells (Fig. 10f).

Next, we grafted samples from patients of the various MB subgroups (Supp. Table 3). Human samples were stained with carboxyfluorescein succinimidyl ester (CFSE) fluorescent vital dye and counterstained with α-mitochondria antibody. As these staining can have variable intensity depending on the samples, we also used another specific human antigen targeting antibody, α-Ku80. As for the murine PDX samples, we observed a small tumor mass forming in the vicinity of the RL for MB SHH (Fig. 10g), MB G3 (Fig. 10h) and MB G4 (Fig. 10i) samples. Thus, placing MB cells from patient samples back to the embryonic context provided by the avian model constitutes appropriate conditions for the cancer cells to survive and manifest their tumoral behavior.

## DISCUSSION

### Modeling MB within its embryonic context of origin

In order to capture yet inscrutable processes occurring at the onset of human MB, we set up an in vivo paradigm to experimentally recapitulate the specific embryonic microenvironment in which MB cells became malignant. We report that human MB cells, when transplanted into the avian embryo cerebellar primordium, survive and form tumors within a few days, benefiting from physiological cerebellar morphogenesis cues, including SLIT. Interestingly, this model allows cells representative of the different MB subgroups to manifest morphological and migratory features reminiscent of their cells of origin, thus recapitulating the heterogeneity of MB. Strikingly concerning MB G4, despite extensive efforts around the world, no animal model compatible with research could be developed so far. Indeed, in mice, intake of grafted MB G4 PDX samples is rare and, when successful, the graft rate is low with a mean latency of 9-12 months^2^. This might be due to the very specific embryonic context in which MB emerges rather than the intrinsic features of this malignancy. Furthermore, once grafted in the chick primitive RL, MB G4 patient samples formed tumor masses in 2 days, thus showing intake and growth capabilities similar to those of other MB subgroups (Fig. 10i). Moreover, for all subgroups, we found that a range of a hundred of tumoral cells was sufficient, for their survival and growth, which represents a 10^3^ to 10^4^ fold reduction, when compared to mice^2^. Additionally, we found that our reductionist eCbOTC model allowed rapid and direct monitoring of MB tumorigenesis (Fig. 1). Compared to previously reported grafts of MB cells within slices of mature murine cerebellum at P8-P10^46^, our model provides to MB cells an environment in which their chick physiological counterparts are concomitantly colonizing the future cerebellum, as in the human pathology^1^.

Despite being a non-mammal vertebrate, the chick holds several key advantages for the study of MB disease within the context of cerebellar development. As in mammals, the cerebellum in birds is folded with folia and lobes^22^ (Supp. Fig. 3). Importantly, the temporal switch between glutamatergic cerebellar cell fates (CN versus GCP) and the SHH-dependent massive EGL expansion have both been acquired by birds. Moreover, DCN and UBCs have been also identified in the chick embryo^11,47^. Thus, cell fates and migration paths of the diverse cerebellar progenitors present in humans are well conserved in birds. In contrast, although offering opportunities for drug screening, studies based on xenograft of MB cells in zebrafish embryo are limited by cerebellar structural and developmental specificities^48,49^. Finally, our chick embryo models complement those based on the murine embryo that is hardly accessible for cell transplantation. Murine embryos were also experimentally manipulated through in utero electroporation of MB oncogenic drivers^50,51^ or the CRISPR/CAS9 machinery to disrupt tumor suppressors^52^. It resulted in the formation of murine tumors and allowed to narrow down the temporal window of MB emergence in early cerebellar development, a finding that was later on confirmed with single-cell transcriptomic analyses. Yet, in these studies, the electroporation of oncogenic drivers generated MB tumors from any targeted cerebellar glial or neuronal progenitor tested, overriding specific lineages of origin and bypassing the first steps of tumor establishment^51^.

### Recapitulating the embryonic context reveals the functional proximity of MB cells with their cells of origin

Interestingly, our paradigm of human MB cells transplantation of different MB subgroups revealed that they manifest functional properties, reflected in diverse morphologies and migration profiles, that match those of their respective cells of origin. First, combining our models with light-sheet and confocal microscopy, we could closely examine how MB cells adapted to the embryonic tissues. Remarkably, we observed that MB cells adopted polarity features of their cells of origin, showing morphological specificities of different cell states within the differentiation trajectory of the lineage: (i) MB G3 cells adopt the amoeboid protrusive shape of RL early progenitors or the unipolar shape with a small process arbored by CN progenitors. (ii) MB SHH cells resemble unipolar and bipolar shapes of more committed GCP progenitors. Finally, (iii) MB G4 cells extend a brush protrusion prototypical of their sub-lineage of origin (UBC) (Fig. 2 and Fig. 5). These morphologies are not only identity traits; they are also instructive of migratory modes and part of the differentiation program of the lineage. Indeed, as already described by Ramon Y Cajal more than a century ago, when they emerge from the RL germinal zone, progenitors are polyhedral and protrusive^53,54^ (Fig. 1a and Supp. Fig. 1a). They rapidly take up a unipolar morphology which enables a saltatory forward movement pulled by a leading process^8,11^. Along this tangential subpial migration, progenitors form chain-like streams and elongate their processes^54,55^. Along their concomitant differentiation and migration, CN maintain this unipolar shape over their concomitant differentiation and migration^8,56^, UBC extend more ramified processes^57^ and GCP become bipolar before initiating their radial migration toward the IGL^54^. These findings suggest that tumor cells might take advantage of these morphological features. Multicellular streaming migration adopted by neuronal progenitors, each pulled by a leading process, has been estimated to range from 0.1 to 1 μm/min^58^ which is considered as a fast migration, often coopted by tumor cells^59^. Various studies have further demonstrated that the rate of progenitor movement correlates with cell shape and particularly the length of the leading process^60–62^. Within the MB cells of all subgroups that we observed, some extended leading processes even reached more than 20μm, in particular for DAOY cells. The surrounding enlarged spatial domain probed by a long leading process is considered as particularly efficient for the perception of gradients of chemoattractants or chemorepulsive cues active in the embryonic cerebellum to guide their physiological counterparts^63^. Altogether, tumors cells might benefit from this fast migration mode and optimized exploratory behavior to adapt to environmental constraints. As observed in Fig. 2a, the dense axon network of the EGL might prevent any wandering of malignant cells in the tissue, favoring their channeling to their destination and their coalescence to form a tumor.

Beyond this, our observations revealed an impressive arsenal of phenotypic features, reflecting a high degree of MB cell plasticity. Such heterogeneity of morphologies is consistent with the conclusions of recent studies. At the time of sampling, cells within the tumor manifested heterogeneous states along the lineage of origin, with MB cells following the developmental trajectory of the lineage of origin and arresting at different levels of the lineage tree^3–5,15,64^. More specifically, two recent studies demonstrated, via spatial transcriptomic, that MB tumors reconstitute a cytoarchitecture resembling that of the cerebellar stratified neuroepithelium, in direct link to the developmental trajectory of tumor cells^65,66^. Our study enhances these observations, showing that tumor cells not only follow the developmental trajectory of origin at the transcriptional level, but also express functional features of their cells of origin, such as morphological polarity, migration pathfinding and topographic organization. Interestingly, these observed functional behaviors might mirror those that occurred in patients. This is supported by the striking similarities of the tumor locations in our model to those reported in MB patients. Hence, MRI imaging demonstrated that MB SHH cells are significantly more likely to develop in the vermis and cerebellar hemispheres, while MB G3 and G4 cells are mostly paraventricular^3,4,25^. In our model, though the injection site was common to all MB subgroups, a few days later, cells had colonized distinct parts of the primitive cerebellum. As in patients, MB SHH settled in the vermis and cerebellar hemisphere underneath the pial surface, where their cells of origin, the GCP, transit and massively expand. MB G3 and G4 migrated ventrally through the future white matter, as do their physiological counterparts, early RL, CN and UBC progenitors (Fig. 4). This is likely to also occur in the human pathology as oncogenic transformation affects these progenitors when they are still migrating to reach their final destination^3–5,15^. Thus, MB cells seem to be able to arrest sooner or later along their differentiation and their migration trajectories.

### Therapeutic interest of targeting polarity and migratory pathways in MB, like SLIT signaling

In an effort to advance our understanding of the primitive molecular dialogues between MB cells and their embryonic microenvironment, we conducted a differential RNAseq analysis, comparing MB cells in culture and in the chick cerebellar primordium. The axon guidance SLIT signaling turned out to be predominantly activated in the embryonic context. SLITs are known to bind several receptors of ROBO and PLEXIN A families, having interactions themselves with other membrane proteins and extracellular ligands that can impact the overall signaling activity (Supp. Fig. 11a). This interaction network makes manipulation of SLIT signaling challenging, for example, due to various receptors compensating the lack of the others^34^. We could, nevertheless, observe that knock-down of ROBO1 and PLXNA1 resulted in the increase of MB G3 cell cohesion and, in mirror, in the decrease of the number of peripherally isolated “pathfinders” cells in the forming primary tumor (Fig. 9 and Supp. Fig. 11b). These effects were significant, although subtle, possibly due to compensation. Interestingly, a recent study identified a significant role for ROBO1 in recurrent Glioblastoma (rGBM)^67^. ROBO1 was reported to participate to rGBM maintenance of stemness, cell proliferation and invasion. CAR-T cells targeting ROBO1 demonstrated a high antitumoral effect in a murine model of orthotopically grafted rGBM cell lines. This CAR-T ROBO1 approach also turned out to be also efficient in the relapsed MB G3 cell line (MB002) in vitro and in orthotopically xenografted adult mice. Our findings suggest that ROBO1-PLXNA1-SLIT2 signaling is activated from the earliest stages of the disease and could be maintained or preferentially reactivated to support the malignant behaviors of MB cells later on during disease progression. This is fully consistent with our analysis of the MB patient cohort showing that tumors expressing high level of SLIT receptors are associated with a bad prognosis and tumor aggressivity (Fig. 10 and Supp. Fig. 9 and 10).

An intriguing possibility is that SLIT-driven emergence of isolated cells at the periphery of the tumor mass initiates an infiltration process (Supp. Fig. 11b). As shown for other de-cohesive guidance cues it might also prepare tumor cells to metastasize^68,69^. Indeed, it was demonstrated in GBM by intravital imaging among other techniques, that the tumoral cells invading the brain parenchyma are isolated and unconnected to the tumor core^70,71^. Such an effect would resemble another physio-pathological function. Indeed, SLIT signaling has been shown to enable neuroblasts committed to olfactory bulb colonization to sneak into the brain parenchyma during normal neurodevelopment^72^. Notably, SLIT signaling is later reactivated by newly generated neuroblasts in the pathological context of stroke where it helps cells to colonize and regenerate a striatal lesion^73^.

Altogether, our avian model offers a great opportunity to decipher the behaviors of MB cells in light of their cells of origin and the embryonic context of initiation. As demonstrated for SLIT signaling, our molecular analysis brings relevant basis for the identification of novel therapeutic targets. Furthermore, our study reveals that the various MB subgroups take advantage of polarity and morphologies changes. And these acquired characteristics seem to be maintained during tumor progression. It would likely be too precarious to investigate theses signaling patterns in patient tumors. With our model and the knowledge of the mechanisms specific to the various lineages of origin, we may be able to identify new therapeutic targets linked to cytoskeleton remodeling that are advantageously exploited in the corresponding MB subgroups.

More generally, various recent studies have proven that driving MB cells toward differentiation constitutes a promising strategy^74,75^. Thus, close comparative analysis of the behaviors of MB cells and of their cells of origin might accelerate the discovery of key molecular levers that would constrain tumoral cells to commit into their differentiated fates.

## MATERIAL AND METHODS

### Chick embryos

Nacked neck embryonated eggs were obtained from a local supplier (EARL Morizeau, Dangers, France). The laying hens’ sanitary status was regularly checked by the supplier according to French laws. Experiments with chick embryos were performed within the first 14 days of gestation, stages that do not require approved protocol by an ethics committee, according to the revised European ethics legislation (2013). Eggs were housed in an 18°C incubator until use. They were then incubated at 38.5 °C in a humidified incubator until the desired developmental stage: HH18 for the graft step (72h of incubation); these embryos were then harvested between HH26 (E5) and HH31 (E7) (early stages) or HH31 and HH40 (E14) (late stages). Embryonic Cerebellum Organotypic Tissue Culture (eCbOTC) were dissected out of HH26 embryos (5 days of incubation).

### Cells lines

Human MB G3 cell lines, HD-MB03 were obtained from DSMZ (DSMZ^©^, ACC740) and grown in RPMI Medium 1640 GlutaMAX™ (Gibco™, 61870044) complemented with 10% Fetal Bovine Serum (FBS, Pan Biotech, P30-2002). SHH MB cell lines, DAOY (ATCC^®^, HTB-186^™^), were a kind gift from P. Mehlen (Centre de Recherche en Cancérologie de Lyon, Lyon, France) and were grown in Dulbecco’s Modified Eagle Medium (DMEM) GlutaMAX™ (Gibco™, 10566016) complemented with 10% FBS. Both previously described media were supplemented with 1 % Penicillin-Streptomycin (PS, Gibco™, 15140122) and 1% Amphotericin B (AB, Sigma-Aldrich^®^, A2942). MB G4 cell line, CHLA-01-MED (CHLA-01) were obtained from ATCC (ATCC^®^, CRL-3021™). These cells are not adherent and were cultured in DMEM-F12 Media (Gibco™, 11330032) complemented with 2% B-27 supplement serum free (50x, Gibco™, 17504044), human recombinant EGF (hrEGF, Gibco™, PHG0314) and human recombinant FGF-2 (hrFGF-2, Gibco™, PHG0026) at a concentration of 20 ng/mL each. All these cell lines were modified to stably express GFP reporter (HD-MB03::GFP, DAOY::GFP, CHLA-01::GFP). Stable expression of GFP was obtained by transduction of HIV1-based lentiviral particles. Lentiviral particles were generated at the lentivectors production facility / SFR BioSciences Gerland - Lyon Sud (UMS3444/US8). Briefly, cells were plated in six well plates (5x10^5^ cells per well) in complete medium. After 2 hours, the medium was replaced with 2 mL medium containing 2 % FBS and 2 mg/mL polybrene (Sigma-Aldrich^®^, TR1003). An hour later, that medium was removed and replaced with 2 mL of medium containing 5x10^6^ IU of lentiviral vector. 16 hours later, the medium was removed, and the cells were rinsed and incubated with complete medium (10 % FBS). Under a fluorescence microscope, nearly 100 % of cells were positive for GFP. Medium from semi-confluent transduced cells showed no capacity to transfer GFP expression to naive control cell lines, indicating that infectious viruses were not produced by the transduced cells. MB G3 cell line, MED8-A::GFP (MED8A^76^ CVCL_M137 modified to express GFP) was cultured in Dulbecco’s Modified Eagle’s medium (DMEM) (Invitrogen) + GlutaMAX-I high glucose medium (Thermo Fisher Scientific) supplemented with 10% fetal bovine serum (FBS) (Merck, #F9665) and 1 % penicillin/streptomycin (P/S) (Merck). Murine MB SHH PTC1MUT tumor cells were obtained from spontaneous medulloblastoma derived from the *Ptch1*^+/-^ mouse model and were cultured as described previously^77^.

### PDX samples

ICN-MB-PDX4 samples (G3 MB) were taken from primary human MB tumors obtained from tumor resections performed at the Children’s Necker Hospital (Paris, France). All human samples were obtained with informed consent of patients or their representatives, and all experimental procedures were performed following guidelines from the Institutional Review Board at Necker Hospital (Paris, France). PDX samples were thawed at 37°C for 2 minutes and resuspended in a PDX medium: Neurobasal medium (Gibco™, 12348017), 2 % B-27 supplement serum free (50x, Gibco™, 17504044), 1 % of N-2 supplement serum free (100X, Gibco™, 17502048), 0.4 % BSA (Bovine Serum Albumin) solution (30 % stock solution, Sigma-Aldrich^®^, A9576), 1 % PS, 1 % L-Glutamine (Gibco™, 11534546). hrEGF (Gibco™, PHG0314) and hrFGF-2 (Gibco™, PHG0026) at 20 ng/mL final concentration each, were added. Samples were resuspended in 1 mL of PDX medium, filtered with a 40 µM membrane (Falcon^®^, 352340), and centrifuged for 10 min at 200 g. They were finally filtered with pluriStrainer Mini 40 µM (Pluriselect, 43-10040-40) and then grafted *in ovo*.

### Patient biopsies

*Sample dissociation*. Samples were thawed at 37°C in a water bath for 2 minutes. They were rinsed 4 times in pre-warmed filtered complete RPMI Medium 1640 GlutaMAX™ (Gibco™, 61870044) complemented with 10 % FBS and 1 % PS. Samples were incubated in 1.5 mL microtube with FBS for 3 min at 37°C and rinsed 3 times with HBSS (Gibco™, 14170088).

Samples were cut in small pieces of 1 mm^3^ in 500 µL of HBSS. The mixture was transferred in a tube containing 1.25 mg/mL of collagenase type IV (Sigma-Aldrich^®^, C5138), 50 mM of CaCl_2_ solution (Sigma-Aldrich^®^, 449709), 0.125 mg/mL of DNase (Sigma-Aldrich^®^, DN25). The mixture was then incubated for 30 min at 37°C and mixed by pipetting up and down every 5 minutes. Trypsin at a final concentration of 0.5 mg/mL was added and the homogenized mix was incubated for 2 min at 37°C. For final filtration, the mix was resuspended in 10 mL of complemented RPMI medium and passed through a 40 µm filter (BD Bioscience, 252340). Samples were rinsed in HBSS and centrifuged for 10 min at 200 g.

*CFSE Labelling.* The cell pellet was transferred in 500 µL HBSS into a 1.5 mL tube and centrifuged for 10 min at 200 g. The pellet was resuspended in 300 µL of warm HBSS containing 7 µM of CFSE reagent (Invitrogen^TM^, C34554) and incubated for 20 min at 37°C. The tube was gently homogenized after 10 min. Finally, cells were incubated for 5 min at 37°C in their complete medium.

The suspension was homogenized, then filtered with pluriStrainer Mini 40 µM (Pluriselect, 43-10040-40) and centrifuged for 5 min at 200 g. These steps were repeated one more time before graft.

### *In Ovo* graft

3 mL of albumin was collected from each egg (HH11-13) with a syringe to allow a small opening of a window in the shell without destroying the embryo. At stage HH18, the chick embryos were grafted with MB cell lines or patient biopsies at the level of the cerebellar primordium, in the RL. MB cells were implanted with a glass capillary connected to a pneumatic PicoPump (World Precision Instruments, PV820) under a fluorescence stereomicroscope. 1 mL PBS supplemented with 1 % Penicillin-Streptomycin (PS, Gibco™, 15140122) and 1 % Amphotericin B (AB, Sigma-Aldrich^®^, A2942) was added on the chorioallantoic membrane of the embryo. Eggs were closed with solvent-free tape and placed back in the humidified incubator until the desired stage.

### Embryonic Cerebellum Organotypic Tissue Culture (eCbOTC) graft

eCbOTC were dissected from HH18 embryos first rhombomere: the neural tissue between the isthmus (rostral part) and the medulla oblongata (caudal part) was kept. Surrounding tissues, especially meninges and cranial nerve buds were removed and an incision was made along the roof plate to spread the explant in an “open book” arrangement. After dissection in HBSS (Gibco™, 14170088), eCbOTC were incubated for at least 2 h (37°C, 5 % C02) in their culture media: Neurobasal™ (Gibco, 12348-017) complemented with 15 mg/mL D-(+)-glucose (Sigma-Aldrich^®^, G7021), 2 % B-27 supplement (50x), 1 % Glutamax™ (Gibco, 35050061), and 1 % PS. MB cells were implanted in the RL or in the pons with a glass capillary connected to a pneumatic PicoPump (World Precision Instruments, PV820) under a fluorescence stereomicroscope. After the graft, explants were set upon porous membranes (Whatman^®^ Nuclepore™ Track-Etched Membranes, WHA10417301), floating on Neurobasal culture media in the central well of a double-well organ culture dish (Falcon©, 353037). Explants were incubated for 5 days (37°C, 5 % C02) and imaged every day with a fluorescence stereomicroscope (Zeiss Axiozoom v16 and camera photometrics coolsnap HQ2, 25x).

### Growth and cohesion assay in eCbOTC

Analysis of mass behavior at the surface of half eCbOTC were made using Fiji software (v2.9.0/1.53t). Measures were limited to Otsu threshold for area, fluorescence intensity and number of particles. For the number of particles, the background was subtracted, and the watershed algorithm was applied to segment big masses and erode function, with 1 passage being applied to remove non clustered isolated cells.

### Immunostaining slices

Grafted chick embryos or grafted explants were fixed for 2 h and 20 min respectively, in 4 % PFA and embedded in 1X PBS, 7.5 % gelatin and 15 % sucrose. Cryostat sections (Leica, CM1950) of 50 µm for full embryos or 30 µm for explants were stored at -20°C. For cell staining, HD-MB03 and DAOY cells were plated on glass coverslips (Dominique Dutscher, 100039N) and incubated for at least 1 h (37°C, 5% CO_2_) in their complete culture media for cells to adhere; CHLA-01 cells were plated on glass slides using Cytospin 4. All cells were fixed 15 min in 4 % PFA.

For all slides, permeabilization and blocking were performed at the same time in PBS, 3 % BSA (Sigma-Aldrich^®^, A7906) and 0.5 % Triton (Sigma-Aldrich^®^, T9284).

Slides were incubated 2 h at room temperature with the following primary antibodies (1/500): anti-α-Mitochondria (Merck Millipore^©^, Clone113 MAB1273), anti-Tubulin-ß-III (BioLegend^®^, BLE80801202), anti-Ki67 (Sigma-Aldrich^®^, SAB5700770), anti-ROBO1 (R&D^®^, AF7118), anti-PLEXNA1 (Abcam^®^ ab23391), anti-SOX2 (Abcam^®^, ab97959), anti-α-SMA (Sigma-Aldrich^®^, A2547).

Slides were incubated with the corresponding secondary antibodies: donkey anti-mouse IgG 555 (Invitrogen^TM^, A31570), donkey anti-rabbit IgG 647 (Invitrogen^TM^, A31573), donkey anti-sheep 555 IgG (Abcam^®^, AB150178). Nuclei were stained with Hoechst 34580 (Invitrogen^TM^, H21486). Slides were mounted in fluoromount mounting media (Sigma-Aldrich^®^, F4680) and imaged either with confocal microscope (Olympus, FV1000, X81) using 4x or 10x for general mass localization and 40x or 60x for cell morphology and cohesion analysis; or with thunder microscope (Leica^®^, DMi8).

### Whole embryonic Cerebellum Organotypic Tissue Culture (eCbOTC) immunostaining and imaging

For whole eCbOTC immunostaining, a protocol was adapted from the whole-mount immunostaining protocol. Briefly, eCbOTC were fixed for 20 min in 4 % PFA. Permeabilization and blocking were performed at the same time in 1X PBS, 3 % BSA (Sigma-Aldrich^®^, A7906) and 0.5 % Triton (Sigma-Aldrich^®^, T9284). eCbOTC were incubated for 2 days with the primary antibody anti-Tubulin-ß-III (BioLegend^®^, BLE80801202) followed by a 2-days incubation with secondary antibody donkey anti-mouse IgG 555 (Invitrogen^TM^, A31570). Partial clearing was achieved by incubation of eCbOTC in successive baths of 50% and 80% glycerol in 1X PBS. eCbOTC were then imaged with confocal microscope (Olympus, FV1000, X81) in chambers filled with 80% glycerol in 1X PBS.

### Tissue clearing, whole-mount immunostaining and Selective Plane Illumination Microscopy (SPIM) imaging of chick embryos

Chick embryos were dissected in HBSS. For early stages embryos (HH26-HH29) the whole head had been preserved whereas for late stages embryos (HH31-HH40), the skin, the forming skull and the eyes, were removed without destroying the meninges, the cortex or cerebellum. In both cases they were and fixed for 2 h in 4 % PFA and rinsed twice in PBS. Late stages embryos were dehydrated in successively increasing concentrations of methanol (Sigma-Aldrich^®^, 179337): 25 %, 50 %, 75 %, 100 %, 100 % (1h per bath) and incubated overnight in H_2_O_2_ 0.2 % methanol (Sigma-Aldrich^®^, H1009). Embryos were then rehydrated following the reverse protocol, inverting the sequence of methanol concentrations. For both early and late stages embryos, blocking and permeabilization were performed in 1X PBS complemented with 10 % DMSO, 0.5 % Triton (100X), 2 % BSA and 100 mM glycine (Roth, 3808.2) for respectively 2 h (early stages embryos) or 4 days (late stages embryos) respectively at room temperature (RT) on a rolling agitator. Primary and secondary antibodies were applied in the blocking solution. Early stages embryos were incubated for 2 nights at RT with the primary antibody and rinsed the following day in 1X PBS complemented with 10 % DMSO, 0.5 % Triton (100X). Then, they were incubated for 2 nights with the secondary antibody and rinsed for 2 days in PBS/DMSO/Triton. Late stages embryos were incubated with the primary antibody for 4 nights at RT on a rolling agitator, then rinsed for 1.5 days, and incubated for 2 nights at RT with the secondary antibody and rinsed for 2 nights. For all different stages of embryos, the following primary antibodies were used (1/500): anti-GFP (Invitrogen^TM^, A-11122), anti-α-mitochondria (Merck Millipore^©^, Clone113 MAB1273). Coupled with the following secondary antibodies (1/500): donkey anti-rabbit IgG 647 (Invitrogen^TM^, A31573), donkey anti-mouse IgG 555 (Invitrogen^TM^, A31570). Dehydration was then performed by increasing concentrations of Ethanol: 50 %, 70 %, 100 %, 100 % Ethanol (1 h or 1 day respectively for early and late stages). Incubation in Ethyl cinnamate (Sigma-Aldrich^®^, 102514985) was performed for 1 day at RT. For SPIM imaging, cleared embryos were glued on holders and immersed in an Ethyl cinnamate bath. Samples were imaged on a light sheet SPIM microscope, UltraMicroscope Blaze (Miltenyi Biotec) equipped with × 1.1 (NA = 0.1), × 4 (NA = 0.35), × 12 (NA = 0.53) objectives. Fluorochromes were excited with 488, 561, or 640 lasers. The following filters were used: 525/20, 595/40, and 680/30 to collect the green, red, and far-red emitted light. Image acquisitions were set using the ImspectorPro software (Miltenyi Biotec). Each channel was acquired sequentially. Images were acquired every 2 μm.

### 3D image reconstruction

3D image reconstruction from SPIM or z-stacks from confocal microscope were treated using Imaris™ software. Optic sections were performed on SPIM images of the embryos to visualize the tumor masses. To facilitate the cartography, 3D reconstruction of the cerebellum was performed using the manual drawing function of Imaris™, outlining the borders of the cerebellum revealed by autofluorescence on serial optic sections. Surface reconstruction was then performed on α-mitochondria staining of the tumor mass.

### Quantification of morphological features

Stained cryosections of eCbOTC grafted with DAOY, HD-MB03 or CHLA-01 cells were imaged as stacks by confocal microscopy (Olympus, FV1000, X81), and reconstituted in 3D using Imaris™ software. Cell surface image treatment was applied on GFP staining to allow the visualization of cells with specific morphological features. At this resolution, cells are particularly tangled at the center of the mass but particular morphologies can still be distinguished. Morphologies were classified according to established descriptions of RL-derived progenitors^20,21^. Cells morphologies from the various cell lines were blind counted and classified for at least 3 different explants for each cell line. At least 80 cells were counted for each cell line.

Stained cryosections of chick embryos grafted with DAOY, HD-MB03 or CHLA-01 cells were imaged as well. As for eCbOTC, the cell surface was reconstituted by Imaris™ software. Cells morphologies from the various cell lines were blind counted and classified for at least 5 different grafted embryos for each cell line. At least 650 cells were counted for each cell line.

### Cartography of tumor sites

3D SPIM image reconstructions of in toto stained embryos were performed on Imaris™ software. Only tumor masses double-stained with both α-mitochondria and α-GFP antibodies were considered to avoid artefacts. The percentages of embryos at early stages (HH26-31, E5-E7) with a tumor mass in the meninges (M), and/or in the RL, and/or in the cerebello-pontine angle (CPA) were calculated. At least 12 embryos were blind analyzed for each cell line. The percentage of embryos at late stages (HH31-40, E7-E14) with a tumor mass in the meninges (M), and/or in vermis (Ve) or cerebellar hemispheres (CH), and/or in the paraventricular zone (PV) were calculated. At least 12 embryos were blind analyzed for each cell line.

### Quantification of tumor mass cohesion and isolated peripheric cells

Imaris™ 3D reconstruction of stained cryosections of chick embryos were treated using the cellular segmentation function of the Imaris™ software according to the average cell diameter of the various cell lines (15 μm for DAOY, 7 μm for CHLA-01 and HD-MB03). The distance of each cell to its 9 nearest neighbors was measured by the software. For better visualization, we applied a spectrum look-up table ranging from 25 μm to 45 μm. We then established a classification of cells isolated from their 9 nearest neighbors from a distance > 3 ad (ad: average diameter of the cell) which were labelled in pink and counted. Cells isolated from their 9 nearest neighbors from a distance < 3 ad were labelled in blue and counted. Various cryosections were analyzed per embryo, at least 3 embryos were analyzed and at least 480 cells were counted for each cell line or condition.

### In Situ Hybridization (ISH)

For in situ hybridization on cryosections, wild type embryos were fixed in 4 % PFA and embedded in 1X PBS, 7.5 % gelatin (VWR^®^, 24360.233) and 15 % sucrose (Sigma-Aldrich^®^, S0389). 25 µm cryosections were cryopreserved at -80°C. The protocol for in situ hybridization for slices protocol was adapted from the Molecular Instruments HCR™ RNA-FISH protocol for generic samples on slides. Briefly, slides were dehydrated with increasing concentration of Ethanol: 25 %, 50 %, 75 % and 100 %, 100 % Ethanol (5 minutes per bath). Slides were then rehydrated with reverse protocol, inverting the sequence of Ethanol concentrations. On the same day, pre-hybridization was performed by incubating slides with probe hybridization buffer for 10 min at 37°C in the hybridization oven. Probes, at the final concentration of 4nM in probe hybridization buffer, were incubated overnight in the oven at 37°C. After a wash in PBS, slides were incubated with increasing concentration of 5x SSCT (Saline-Sodium Citrate, Sigma-Aldrich^®^, S6639 with 0.1 % Tween 20X, Electron Microscopy Sciences, 25564) diluted in probe wash buffer: 25 %, 50 %, 75 % and 100 % SSCT (15 min, 37°C in the oven). Hairpin solution was heated at 95°C for 5 min and left to cool down at RT. After a 30 min pre-amplification, the hairpin solution at a final concentration of 60 nM in the amplification buffer was applied overnight at RT. Slices were mounted in fluoromount mounting media (Sigma-Aldrich^®^, F4680)

In toto in situ hybridizations were performed according to previously published protocol^38^. *SLIT2* (Molecular Instruments, NM_001267075.2, Amplifier-fluorochrome: B2-Alexa546) or *PAX6* (Molecular Instruments, NM_205066.2, Amplifier-fluorochrome: B2-Alexa546) probes were designed on chick sequence and purchased from Molecular instruments.

### Plasmids, siRNAs, cell transfection

Control siRNA (Sigma-Aldrich^®^, SIC001), PLXNA1 siRNA (Sigma-Aldrich^®^, SASI_Hs0010385) and ROBO1 siRNA (Sigma-Aldrich^®^, SASI_Hs0100129946) were used at a concentration of 50 nM. For siRNA and plasmids transfections, cells were transfected with JetPrime (PolyPlus, 101000015) according to the manufacturer’s guidelines.

### Hanging drop assay and cohesion quantification

Native or siRNA transfected HD-MB03 cells were resuspended at a concentration of 5.10^6^ cells/mL and cultured in 25 µL hanging drops on the inside lid of a culture dish. Cells were resuspended in complete culture medium as positive aggregation control or in RPMI 1% FBS media complemented with 2 µg/mL recombinant proteins (rhSLIT2 N; rhSLIT2 C). Plates were incubated for 48 h at 37°C, 5 % CO_2_.

The hanging drops were imaged under a fluorescence stereomicroscope (Zeiss Axiozoom v16 and camera photometrics coolsnap HQ2).

Images were analyzed with ImageJ software. The outline of the aggregate was manually selected and the percentage of area occupied by fluorescent cells was calculated by the software.

### Cell proliferation and apoptosis assay

HD-MB03 cells were plated at a density of 5.10^4^ cells per glass coverslip, in their complete culture medium and incubated for 24 h at 37°C, 5 % CO_2_. Cells were treated with RPMI, 1 % FBS, 1 %PS, 1 % AB medium complemented with 2 µg/mL of SLIT2 C human recombinant protein (rhSLIT2 C, Biotechne^®^, 9379-SL-050) or 2 µg/mL SLIT2 N human recombinant protein (rhSLIT2 N, Biotechne^®^, 8616-SL-050) or both.

For proliferation assays, RPMI 20 % and 1 % FBS media were used as positive and negative controls respectively. For apoptosis assays, cells were treated with Staurosporin (Sigma-Aldrich^®^, S6942) at a final concentration of 0.46 µg/mL as a positive control or with 10% FBS RPMI medium as a negative control. Staurosporin treatment was stopped 4 h later and cells were collected. After 48 h in culture (37°C, 5 % CO_2_), cells were fixed for 20 minutes in 4 % PFA. Permeabilization and blocking were performed at the same time in 1X PBS, 3 % BSA (Sigma-Aldrich^®^, A7906) and 0.5 % Triton (100X, Sigma-Aldrich^®^, T9284). To assess cell proliferation, anti-Ki67 (1/200; Sigma-Aldrich^®^, SAB5700770) primary antibody was used in combination with FP547 anti-rabbit IgG (1/500, Interchim^®^, FP-SB5110) secondary antibody. To assess cell apoptosis, anti-cleaved caspase 3 (1/200, cell signaling, #9661) was used in combination with FP547 anti-rabbit IgG (1/500, Interchim^®^, FP-SB5110) secondary antibody. Nuclei were stained with Hoechst 34580 (1/1000, Invitrogen^TM^, H21486). Cells were imaged with a confocal microscope (Olympus, FV1000, X81) using a 10X objective. Quantifications were made on ImageJ software using “Analyze particles” plugin. A ratio of Ki67 or active-caspase 3 positive cells over the total number of GFP positive cells was calculated.

### In vitro cell growth assay

For in vitro cell growth assay between 2x10^5^ and 2.5x10^5^ HD-MB03, DAOY or CHLA-01 were plated in 10 mL of their respective culture media in small flasks. Each day, for 96 h after seeding, the number of cells in a flask was counted using a Counting chamber, (Fast Read® 102, Kova International, BVS100H). Cells were stained with Trypan blue to evaluate viability.

### Bulk-RNA seq Analysis

HD-MB03 cells were grafted in the future cerebellum of HH18 (E3) chick embryos and the tumor masses were micro-dissected at HH28 (E6) under a stereomicroscope. Tumor masses from hundreds of embryos were pooled to constitute each of the 3 cell batches analyzed independently. GFP positive cells were indeed sorted from the microdissected tumors using CellenOne device (Cellenion, France). “NA” isolation, RNA-Seq processing and data analysis were performed at the ProfileXpert core facility (Lyon, France). The total RNA from 3 independent experiments (1 pool of cells post-graft and 1 pellet of HD-MB03 cells in culture for each experiment) was extracted with RNeasy micro kit (Qiagen) and the quality was checked with a Bioanalyzer 2100 (Agilent, RIN >8.0). Ribosomic RNA was depleted with Ribo-ZeroÔ Gold Kit (Epicentre). RNA pre-amplification was performed with 500 pg RNA with Ovation RNAseq system Kit (Nugen). RNA-seq libraries were prepared with 100 ng ds-cDNA with Ovation ultralow library system kit (Nugen) and sequenced using the HiSeq 2500 platform (Illumina, 50 bp single read).

After, the trimming (Trimmomatic v0.39) and quality check (FASTQC v0.11.9) reads were mapped against the human genome (GRCh38) using STAR v2.7.0a. Transcript quantifications were performed with HTSeq-counts (v0.13.5). Data are available on GEO platform (GSE286482, token for reviewers ehupmcmintipdsf). All further bio-informatic analyses were made using R (v 4.4.1). Differential analysis was performed using DESeq2 package (v1.26.0) with a non-blinded log transformation. Batch effect was corrected using Limma package (v3.42.2) and checked with PCA function of DeSeq2 package. All normalized counts were used for comparison with RNA-Seq signatures extracted from previously published papers. Briefly, a list of genes associated with specified signatures was extracted from literature and their expression in our gene set was plotted with pheatmap package (v1.0.12) with a row scaling. Genes with adjusted Benjamini-Hochberg (BH) corrected p-value (adj p-val < 0.05) were used for further enrichment analysis. Gene set enrichment analysis (GSEA) was performed on Reactome Database using ReactomePA package (v1.48.0) and visualized with enrichplot (v1.24.2). Gene sets were considered significantly enriched if their BH adjusted p-value was < 0.05 and when they contained between 5 and 500 genes.

Over representation analysis (ORA) was carried out on genes with absolute log2FoldChange (abs log2Fc) >1.5 using the Gprofiler website (https://biit.cs.ut.ee/gprofiler/gost), human genome: GRCh38.p13) applied to Gene Ontology Biological Process Database (GO-BP, release from 04.12.2024). 176 gene sets were identified as significantly over-represented (BH fdr adj p-val < 0.05) and further used to draw an enrichment map in Cytoscape (v3.10.2, Enrichment Map plugin) with the following cutoffs: overlap coefficient < 0.5, fdr q-value < 0.05 and p-value < 0.001.

### RT-qPCR

For qRT-PCR analysis, total RNA was extracted from different cell lines using the Nucleospin RNAII kit (Macherey-Nagel^®^, NZ74095550). 1 μg of total RNA was reverse-transcribed using the iScript cDNA Synthesis Kit (BioRad, 1708890). qRT-PCR was performed using the LightCycler480 SYBR-Green I Master1 kit (Roche^®^ Life Science, 04707516001) and the CFX Connect Real-Time PCR Detection System (BioRad). The following list of primers were used in the study:

GAPDH: PrimerPCR SYBR Green Assay: GAPDH, Human; qHsaCED0038674

ROBO1: PrimerPCR SYBR Green Assay: ROBO1, Human; qHsaCID0007521

ROBO2: PrimerPCR SYBR Green Assay: ROBO2, Human; qHsaCID0014976

PLXNA1: PrimerPCR SYBR Green Assay: PLXNA1, Human; qHsaCID0011812

### Analysis of published scRNA-Seq data

Previously published sc-RNASeq data of developing cerebellum from Aldinger and al.^42^, were downloaded from authors’ website (https://cells.ucsc.edu/?ds=cbl-dev). Normalized count matrix was extracted. Cells from clusters known to derive from the RL (“02-RL”, “03-GCP”, “04-GN”, “05-eCN/UBC”) were extracted using Seurat (v4.3.0) and updated to Seurat (v5.0.1). Using the publications variable features, PCA was applied, followed by UMAP calculation using the first 15 principal components. Level of expression of genes of interest was plotted on the newly generated map.

### Analysis of published cell lines RNAseq dataset

Bulk RNA-seq expression data for medulloblastoma cell lines were retrieved from the Gene Expression Omnibus (GEO) database. For DAOY cells, samples from Badenetti et al,.^78^ (GSE273794/GSM8437081,GSM8437082,GSM8437083,GSM8437084) and Yuan et al,.^79^(GSE188813/GSM5690436,GSM5690437,GSM5690438), for HD-MB03, samples from Masurkar et al,.^80^ (GSE198463/GSM5949180,GSE174620/GSM5320663) for MED8A, samples from Zou et al,.^81^ (GSE194217/GSM5830910,GSM5830911,GSM5830912) and for CHLA-01-MED, samples from Yuan et al,.^79^ (GSE188813/GSM5690448,GSM5690449,GSM5690450) were analyzed. We verified sample annotations to ensure that only non-modified cell lines were included. When available, we selected unbothered parental cell lines, and there as only 1 study (GSE188813) for which we used control vector expressing cell lines. Raw count tables were downloaded directly from GEO, merged by gene identifiers, and normalized to counts per million (CPM) using custom R scripts (R v4.4.1) to enable gene expression comparisons across cell lines. Graphical representations were generated using ggplot2 (v3.5.1).

### Analysis of published patients’ expression and survival dataset

Published survival data analyses were performed using R (v4.4.1). Cavalli and al. expression data were extracted from the GEO platform (GSE85217) and normalized using normalizeBetweenArrays function from Limma package. Survival analysis focused on MB G3 patients stratified according to the combined expression of both ROBO1 and PLXNA1 receptors as follow:

- Expression(ROBO1) + Expression(PLXNA1) > Mean(ROBO1) + Mean(PLXNA1) -> “high”
- Expression(ROBO1) + Expression(PLXNA1) < Mean(ROBO1) + Mean(PLXNA1) -> “low”

Survival analysis and Kaplan Meyer curves were generated using R package survival (v3.6-4) and survminer (v0.4.9).

### Analysis of in-house cohort RNA-seq data (primary tumors and cell lines)

RNAseq expression analysis was performed on a cohort of 341 primary medulloblastoma, collected by Ayrault Lab and as well as 16 medulloblastoma cell lines prepared by Ayrault Lab, respectively, as previously described in details (Bernardi *et al,.* under revision). Briefly, RNAseq raw data were firstly processed using an in-house pipeline developed at the Institut Curie Bioinformatics Core Facility, following standard analysis protocols in the field and available at https://github.com/bioinfo-pf-curie/RNA-seq. Then raw counts were normalized using the vst (Variance Stabilizing Transformation) method from Bioconductor DESeq2 package^82^. The accession number for the RNAseq raw dataset will be available in EGA repository once the Ayrault’s paper will be published.

Survival analysis focused on MB G3 patients stratified according to the ROBO1 and PLXNA1 expression separetely as well as the combined expression of both receptors as previously described:

- Expression(ROBO1) + Expression(PLXNA1) > Mean(ROBO1) + Mean(PLXNA1) -> “high”
- Expression(ROBO1) + Expression(PLXNA1) < Mean(ROBO1) + Mean(PLXNA1) -> “low”

Survival analysis and Kaplan Meyer curves were generated using R packages survival (v3.6-4) and survminer (v0.4.9) and pvalue was computed using log-rank test. Barplots, boxplots and Pearson correlation analysis were generated using R packages ggplot2 (v.3.5.2) and ggpubr(v.0.6.0), respectively.

### Analysis of in-house cohort proteome data

Samples were profiled by Mass Spectrometry technique as previously described in details (Bernardi *et al,.* under revision). Briefly, the mass spectrometer was operated in data-independent analysis (DIA) mode for peptide acquisition. Raw data files were analyzed using Spectronaut 19 with the spectral library build inhouse and further processed and normalized using myProMS v3.10 (https://github.com/bioinfo-pf-curie/myproms^83^. The accession number for the Proteome raw dataset will be available in PRIDE repository once the Ayrault’s paper will be published. Boxplots were generated using R package ggplot2 (v.3.5.2).

### SLIT2 interaction map

To generate the map of SLIT2’s principal interactors, STRING online database (v12.0) was used. Network was extracted and mapped on Cytoscape (v3.10.2) to manually add the PLXNAs family which have recently been characterized as SLIT receptors^33^.

### Quantification and statistical analysis

Statistical analyses were carried out using Prism software (v10.3.1). All used statistical tests and statistical confidence levels are mentioned in figure legends.

## ACKNOWLEDGEMENTS

We wish to thank the ProfilExpert core facility (Lyon, France) and especially Joël Lachuer and Severine Croze for fruitful assistance in RNASeq data analysis. We thank Clément Emptaz, Anthony Terra from NeuroBioTec, CRB HCL (BB-0033-00046) (Lyon, France) for the assignment of frozen human medulloblastoma samples. We thank Gianluigi Atzeni, Guilhem Tourniaire from Cellenion (Lyon, France) for the sorting of GFP positive cells. We thank Denis Ressnikoff, Bruno Chapuis and the CIQLE plateform (Lyon, France), Homaira Nawabi, Stéphane Belin, Noémie Vilallongue (GIN, Grenoble, France) for technical advice and discussions. We thank Olivier Imbaud and Benjamin Villalard for their help in bioinformatics and Edmund Derrington for lentiviral infections, Manuela d’Allessandro, Maé Chapuis, Aya Chamaa and Léa Mayoute for technical help. We thank Célio Pouponnot and Ludovic Telley for scientific discussions. We also thank the React4kids French national network in fundamental research in pediatric oncology for stimulating discussions. We would like to particularly thank Loraine Jarrosson and Clélia Costechareyre (Oncofactory, Lyon, France) for helpful discussions and Hannah Gearing for manuscript correction.

This work has been supported by Equipe labelisée Ligue contre le cancer (V.C.) Fondation ARC pour la recherche sur le cancer Programmes Labellisés N° ARCPGA12021020003088_3559 (V.C.), la Ligue contre le Cancer N°AAPEAC2019L.CCS/TD (S.T-D) and N°AAPEAC2023.LCC/STD (S.T-D). This work was conducted within the framework of the LABEX CORTEX, LABEX DevWeCAN of Université de Lyon and South-ROCK INCa-Cancer_18695, and React4kids. MM PhD fellowship was financed by “Une nuit pour 2500 voix”, “Enfants Cancer Santé” associations.

## AUTHOR CONTRIBUTIONS

S.T.D. designed and oversaw the study. C.D.B., M.M., F.M., K.T., C.I. and M.S. made experiments in the avian embryo model. M.M. performed the bioinformatic analyses of RNA-seq datasets from the avian embryo model, and from the publicly available data with support from J.G. and J.T.D, under the supervision of S.T.D. T.F. and C.F.C. contributed to patients’ samples preparation and collection. J.F. set up imaging pipeline. V.C. contributed to analysis results, interpretation of findings. O.A and D.M. provided input to the study and biological samples. S.T.D. wrote the manuscript with the contribution of M.M. V.C. reviewed and edited the manuscript.

## DATA AVAILABILITY

Source data are provided with this paper. Information and data provided in the present manuscript, figures, supplementary information and source data is sufficient to assess whether the study claims are supported by the data. Raw lightsheet and confocal microscopy files generated in the study are available upon simple request. Raw sequencing data from MB implanted in the avian chick embryo have been deposited in GEO database under the accession number GSE286482 (token for reviewers ehupmcmintipdsf)).

