## Supplemental Figures for "Lineage of origin-specific developmental programs drive the behaviors of malignant cells in an avian embryo model of human Medulloblastoma subgroups"

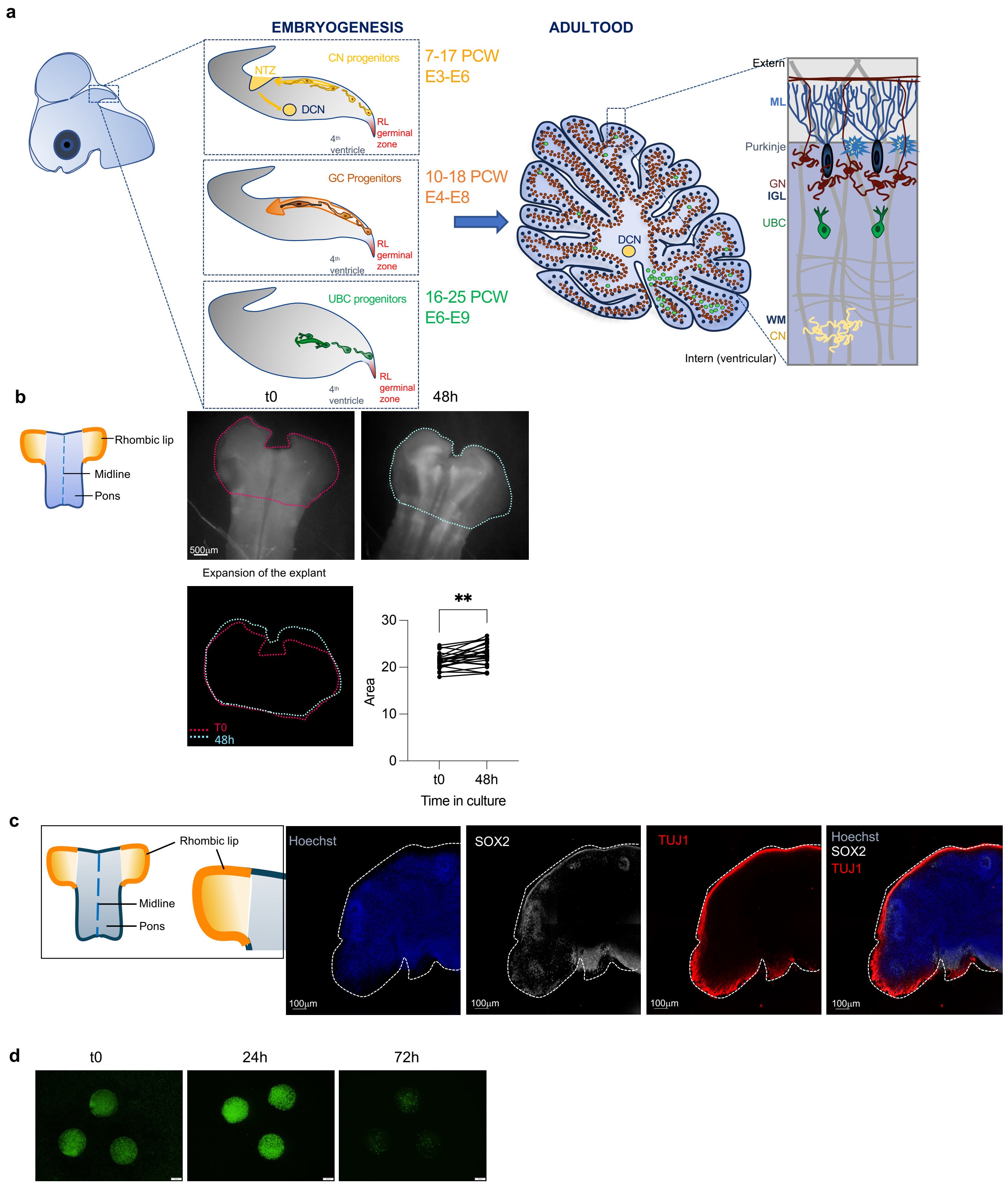


Supp. Figure 1

**a**

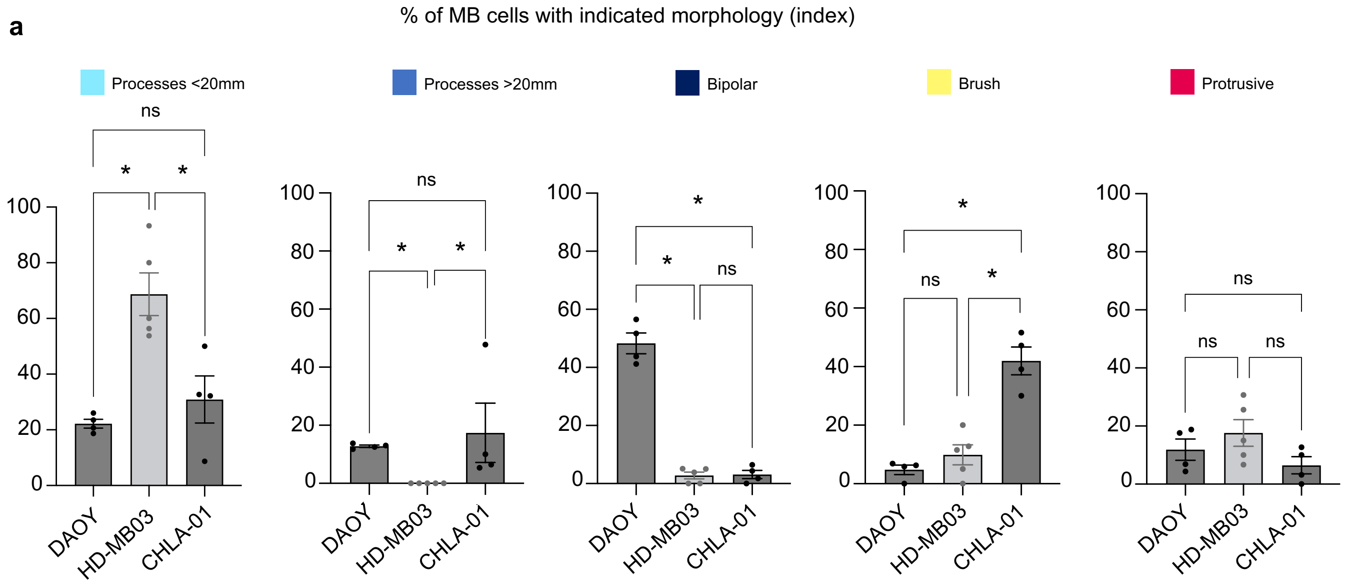


**Supp. Figure 2**

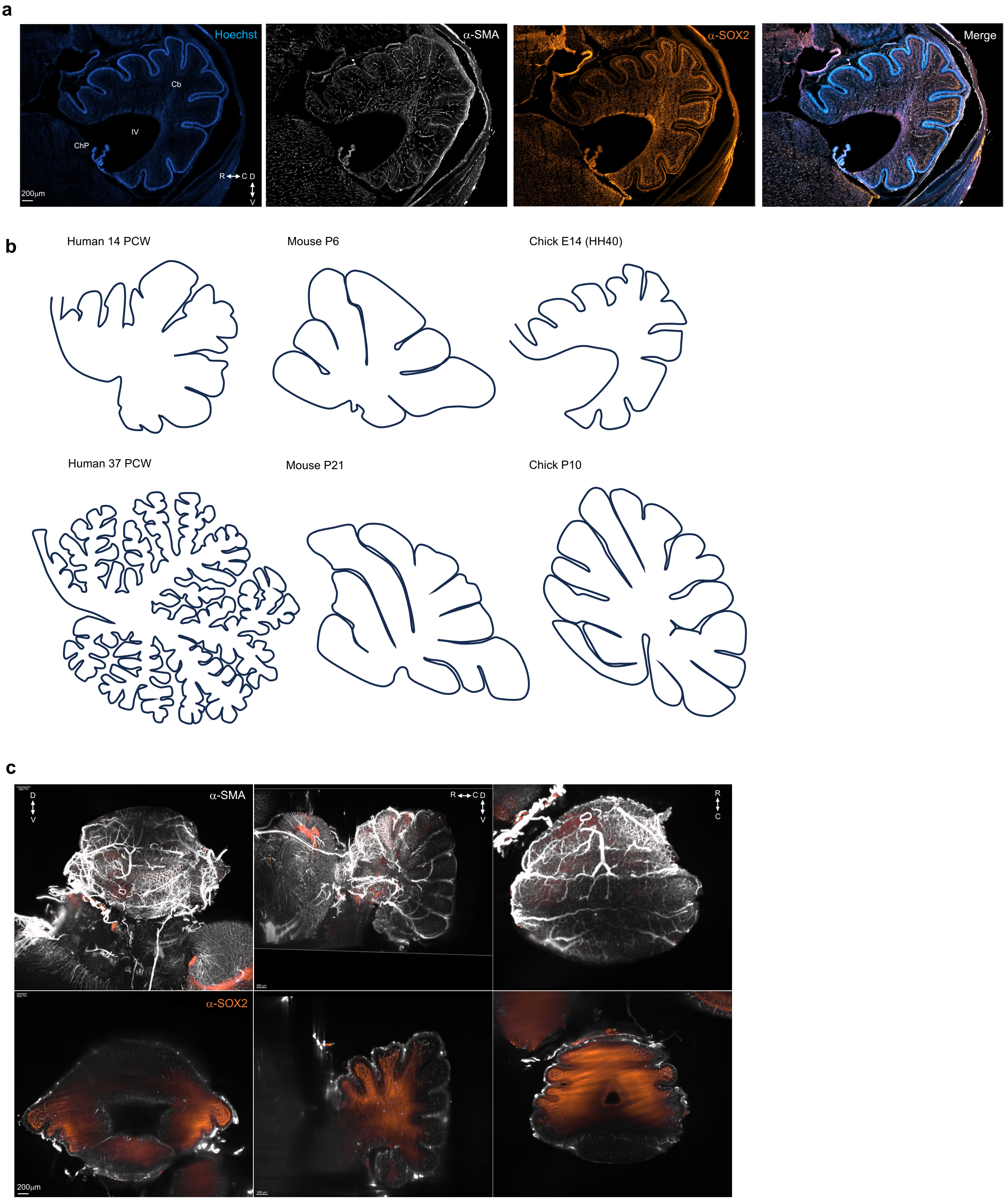


**Supp. Figure 3**

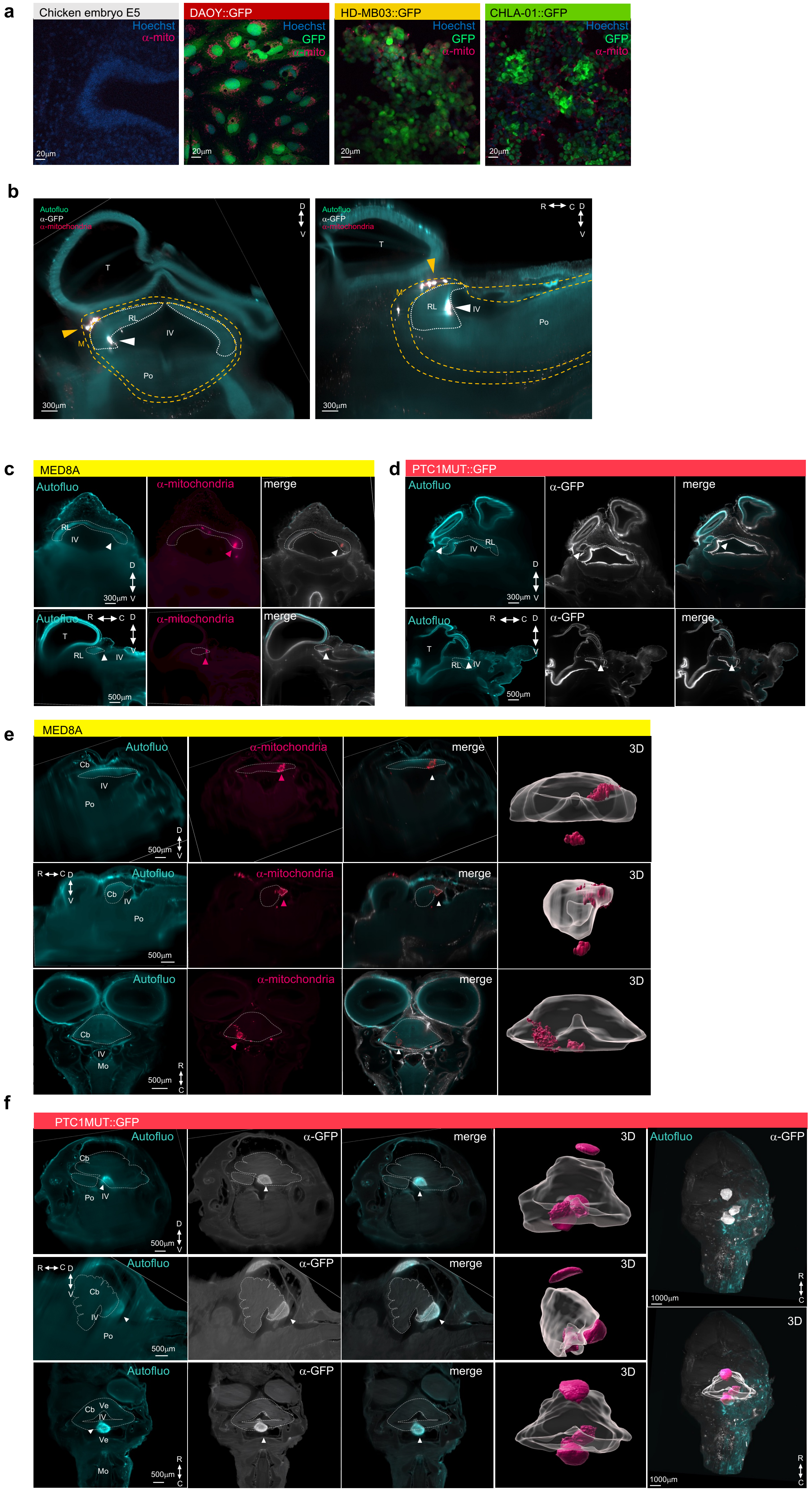


Supp. Figure 4

a

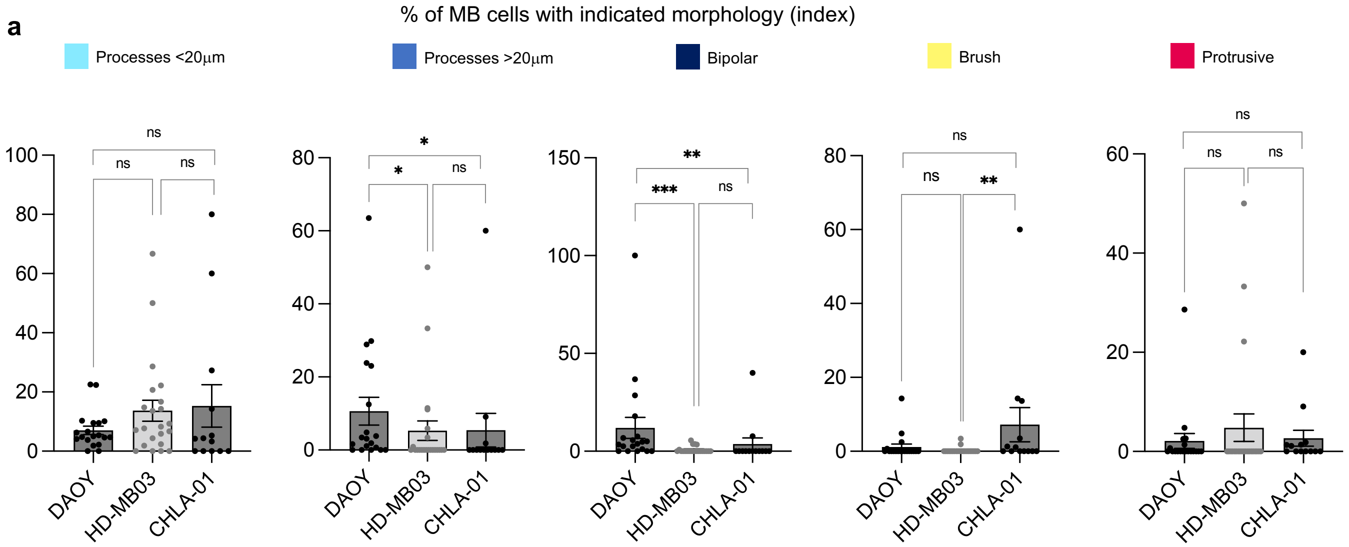


Supp. Figure 5

a

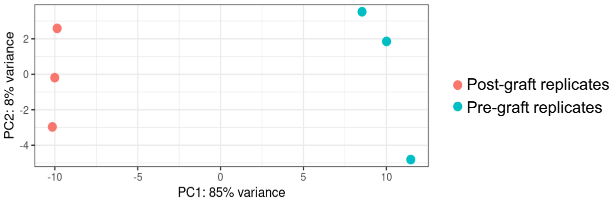


b

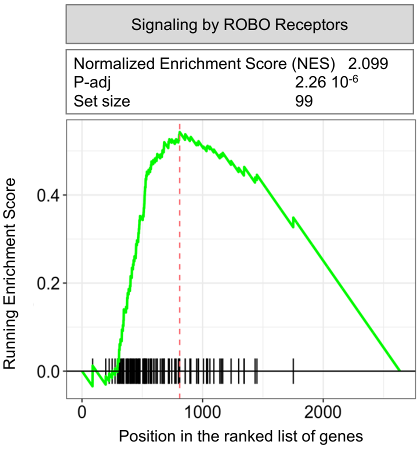


Supp. Figure 6

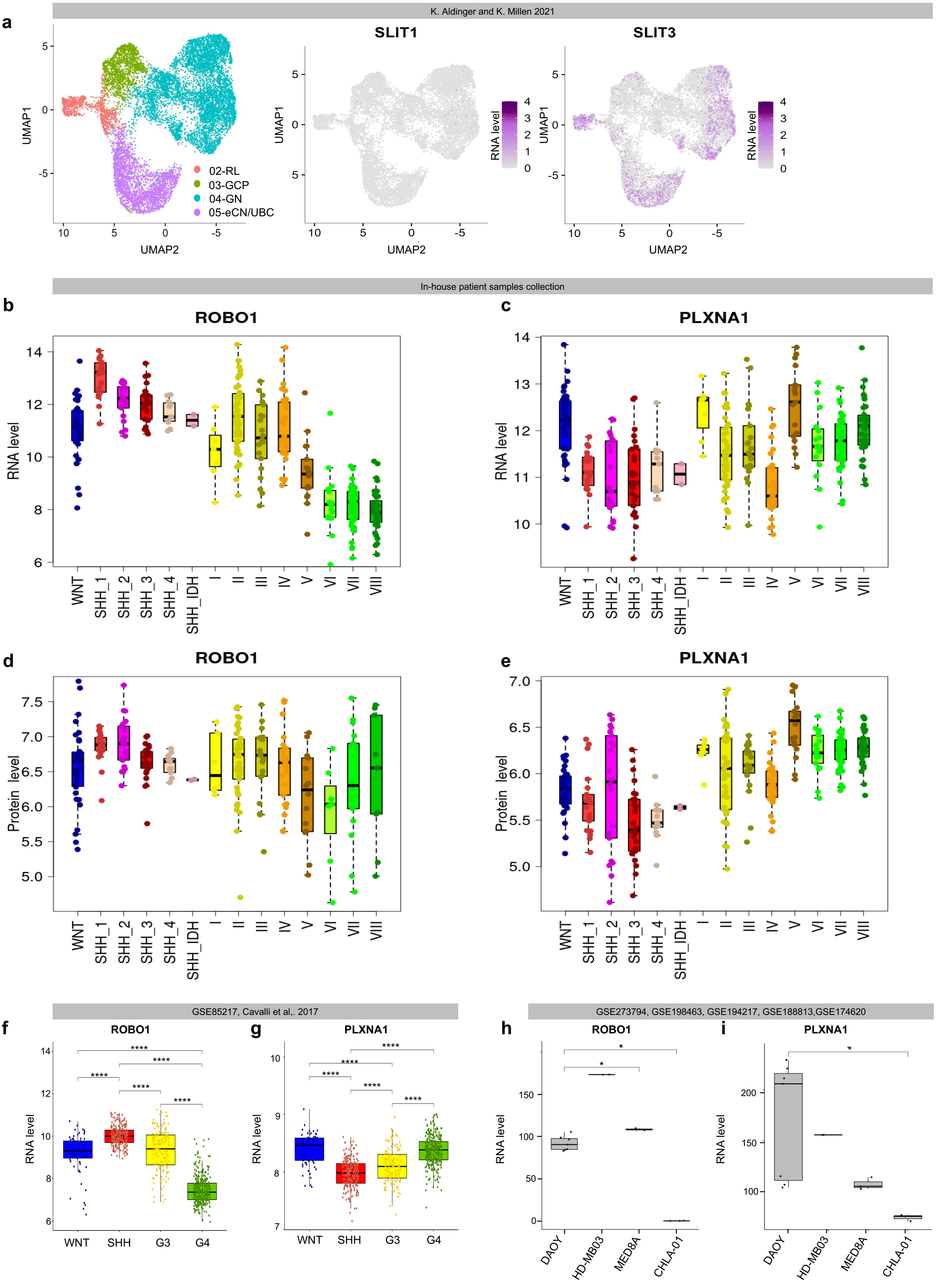


Supp. Figure 7

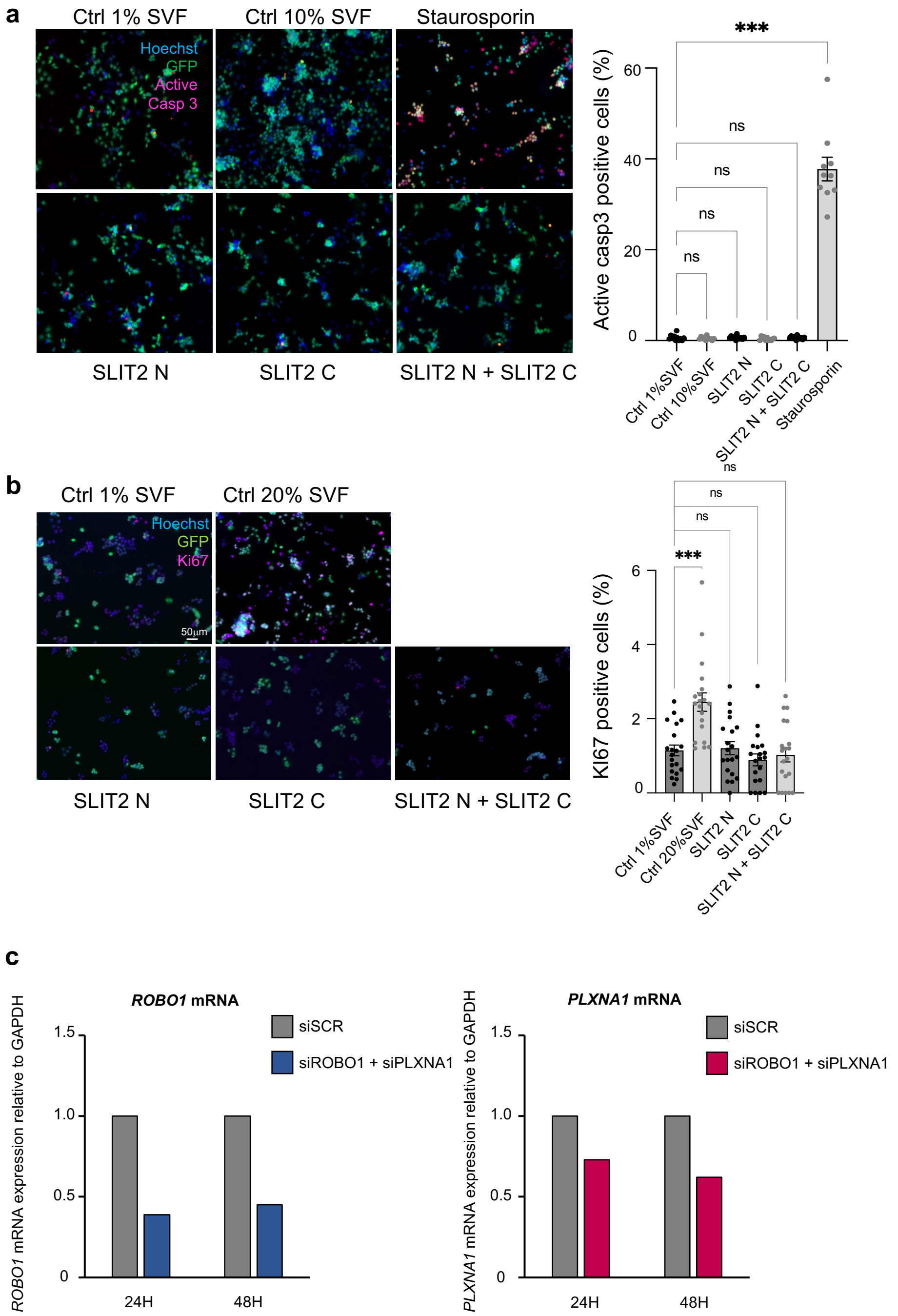


**Supp. Figure 8**

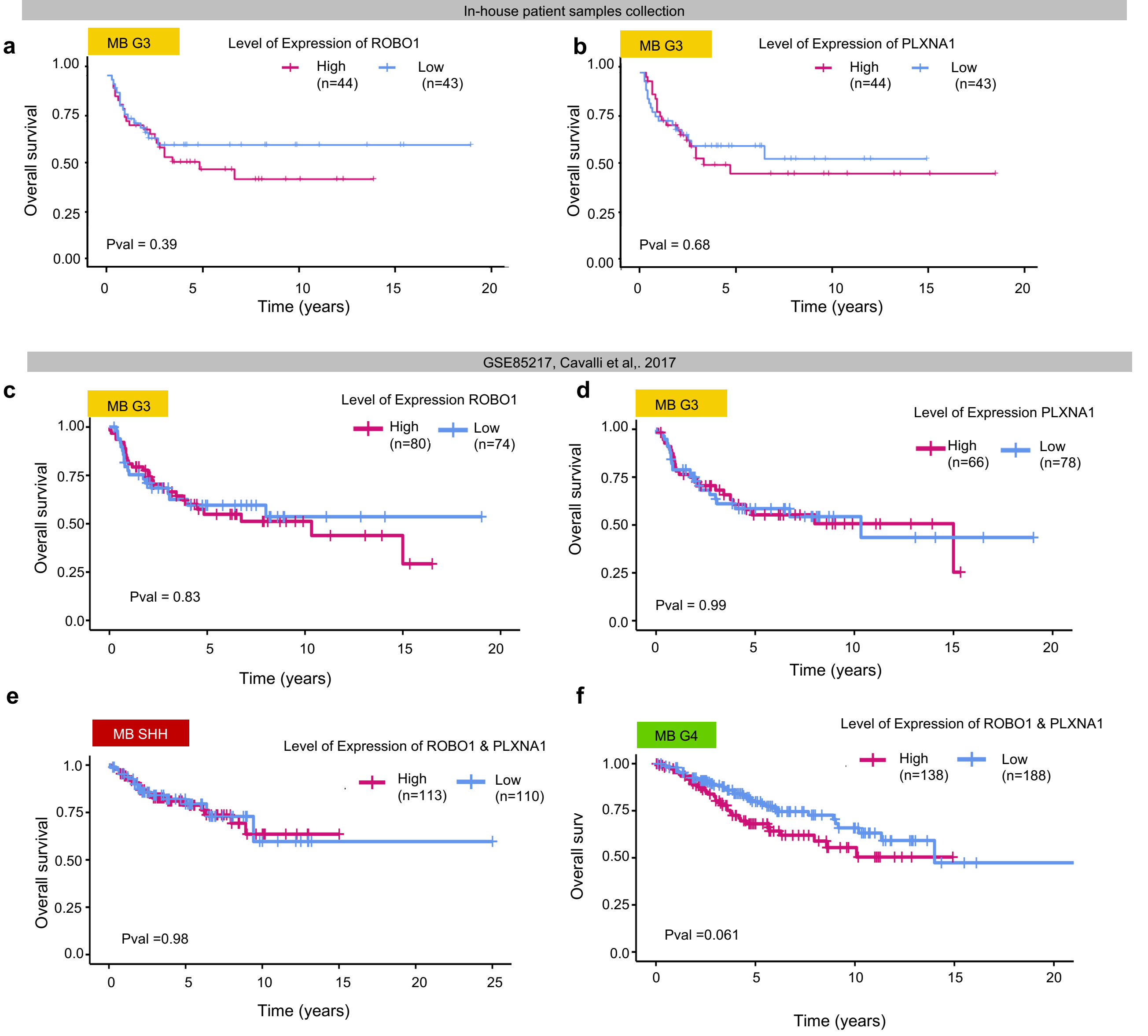


**Supp. Figure 9**

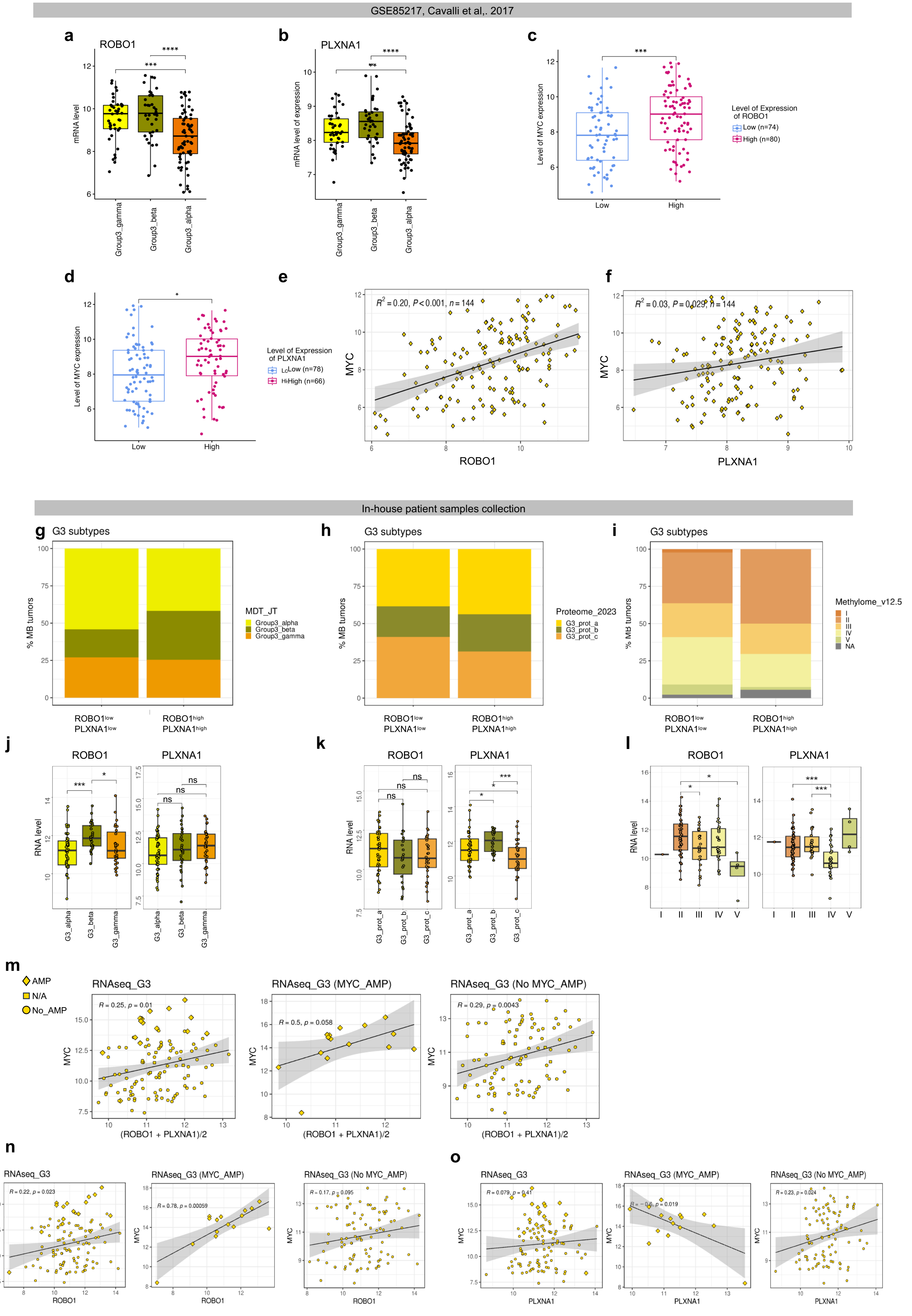


Supp. Figure 10

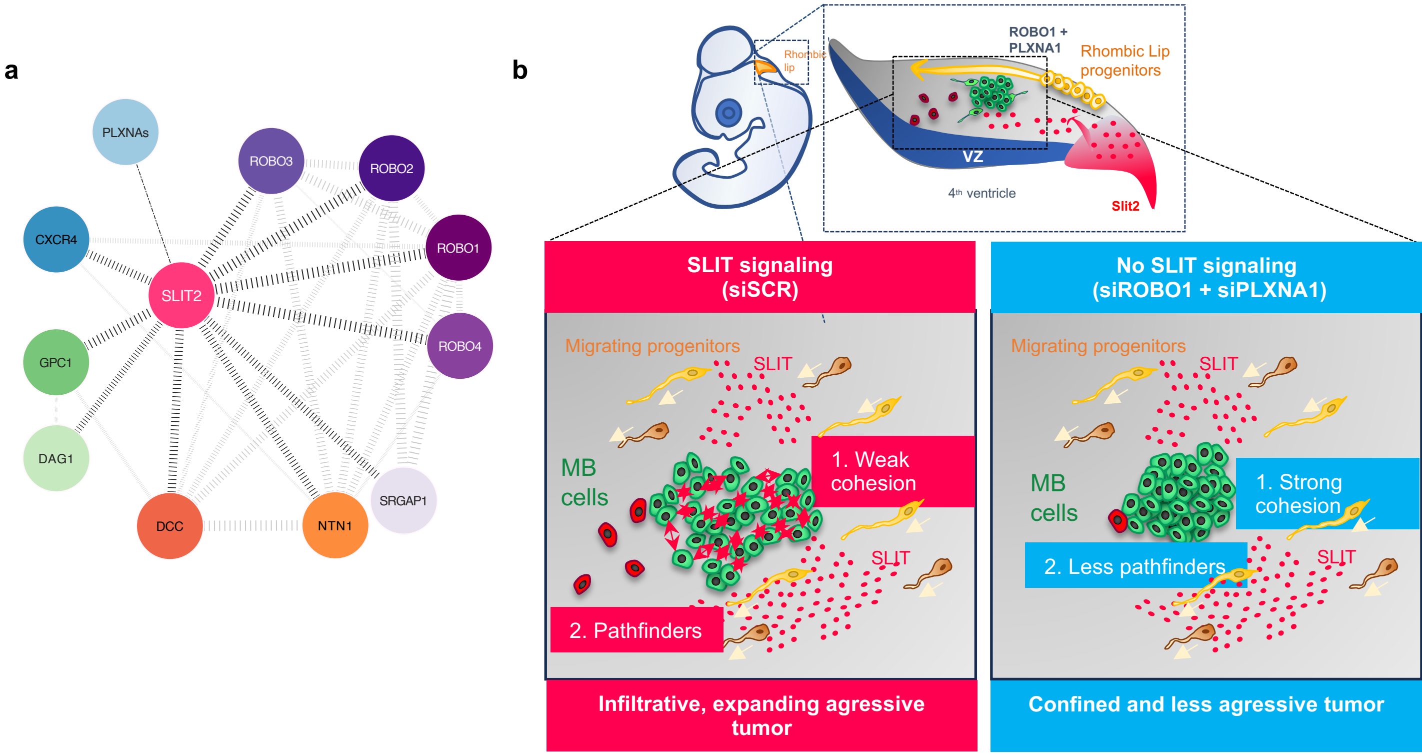


Supp. Figure 11

**SUPPLEMENTAL FIGURES LEGENDS**

**Suppl. Figure 1. eCbOTC model partially recapitulates the early steps of cerebellum development. a.** Schematic representation of the successive waves of RL-derived progenitors emerging from the germinal zone during embryogenesis (left panel). The approximate stages of emergence in human (PCW, post conception week) and in the chick embryo (E, Embryonic day) are indicated on the right. A schematic representation of a sagittal section of a mature chick cerebellum is represented (right panel) with a zoom on a portion of cerebellar stratified neuroepithelium with the various neuronal populations. CN: Cerebellar Nuclei, DCN, Deep Cerebellar Nuclei, NTZ: Nuclear Transitory Zone, GC: Granular Cell, UBC: Unipolar Brush Cell, WM: White Matter, ML: Molecular Layer, GN: Granular Neurons, IGL: Inner Granular Layer. **b.** Representative stereomicroscopic images of the embryonic cerebellar whole tissue explant (eCbOTC) cultured in "open-book" as indicated on the left. The observation and measurement of the area of the explant were performed at t0 (in pink dotted lines) and after 48h (in blue dotted lines) and a significant growth was observed (n=11). A Wilcoxon matched-paired test was performed, $^{**}p \leq 0.01$. **c.** Macroscopic observations of in toto staining of ungrafted eCbOTC with Hoechst (nuclei), SOX2 (progenitors), b-Tubulin III (TUJ1) (neuronal projections) antibodies. **d.** Spots of HD-MB03::GFP cellular suspension were grown directly on the micropore membrane in contact with the culture medium. GFP signal faded after 72h indicating that the cells do not survive.

**Suppl. Figure 2. Once grafted in eCBOTC, MB cells integrate into the tissue and adopt morphologies reminiscent of their lineage of origin. a.** Statistics corresponding to Fig. 2f. , * $p \leq 0.05$. The Mann-Whitney tests were performed to compare 2 cell lines at a time for each morphological feature.

**Suppl. Figure 3. Early developing cerebellum in the chick embryo displays cytoarchitectural similarities with that of human and mice embryos. a.** Representative images of cryosections of E14 (HH40) ungrafted chick embryo stained with Hoechst (nuclei), a-SMA (blood vessels) and SOX2 (progenitors) antibodies. **b.** Schematic drawing of cerebellar foliation in equivalent developmental stages comparing human, murine and chick embryo. Adapted from Haldipur *et al,.* 2022. **c.** SPIM imaging of the cerebellum of a chick embryo at E12 and stained with a-SMA and SOX2 antibodies. The embryo is observed from the front (left), from the side (middle) and from the back (right) and the corresponding optic sections (lower panel).

**Suppl. Figure 4. The topography of the tumors formed by MB SHH PTC1MUT and MB G3 MED8A cells reflects those observed in patients. a.** Representative images of a cryosection of an ungrafted chick embryo (E5) and DAOY::GFP, HD-B03::GFP and CHLA-01::GFP cells in culture stained with Hoechst and the anti-mitochondria antibody which specifically stains human antigens. **b.** Representative coronal (left) and sagittal (right) optic sections of a chick embryo 72h post graft (CHLA-01::GFP) stained with anti-GFP and anti-mitochondria antibodies. Autofluorescence (autofluo) allows the visualization of structures. The whole embryo was imaged with SPIM. The RL is delineated by a white line and the tumor mass in the RL is indicated by a white arrow head. Meninges (M) along the RL and the pons are delineated by yellow lines and the tumor masses in the M are indicated by a yellow arrow head. **c.** Representative coronal (upper panel) and sagittal (lower panel) optic sections of a chick embryo 72h post graft of MB G3 MED8A cells in toto stained with an anti-mitochondria antibody targeting human cells. Autofluorescence (autofluo) allows the visualization of structures. The whole embryo was imaged with SPIM. Arrowheads point to the tumoral masses in the RL (RL) delimited by white doted lines. **d.** Representative coronal (upper panel) and sagittal (lower panel) optic sections of a chick embryo 72h post graft of murine MB SHH PTC1MUT::GFP cells. In this case of murine cells, only GFP and autofluorescence allows the visualization of the grafted cells. **e.** Representative coronal (upper panel), sagittal (middle panel) and transverse (lower panel) optic sections of a chick embryo 7 days after the graft of MB G3 MED8A cells. The embryo was in toto stained with an anti-mitochondria antibody targeting human cells. 3D reconstruction was performed with Imaris software to better visualize the topography of the tumor. **f.** Representative coronal (upper panel), sagittal (middle panel) and transverse (lower panel) optic sections of a chick embryo 10 days after the graft of murine MB SHH PTC1MUT::GFP cells. The embryo was in toto stained with an anti-GFP antibody as anti-mitochondria antibody does not stain murine cells. For e and f, autofluorescence (autofluo) allows the visualization of structures. The whole embryo was imaged with SPIM. Arrowheads point to the tumoral masses in the paraventricular zone (e) or the cerebellar hemispheres (f) delimited by white dotted lines. 3D reconstruction was performed with Imaris software to better visualize the topography of the tumor. The cerebellum outlines are represented in white and the tumor mass surface reconstruction is in pink. RL: RL, CPA: Cerebello-Pontine Angle, IV: fourth ventricle, Po: Pons, T: tectum, Cb: cerebellum, D: dorsal, V: ventral, R: rostral, C: caudal, A: anterior, P: posterior.

**Suppl. Figure 5. At the microscopic level, the behavior of grafted MB cells in the primitive cerebellum reflects that of their lineage of origin. a.** Statistics corresponding to Fig. 5f. Mann-Whitney tests were performed to compare 2 cell lines at a time for each morphological feature. Error bars indicate SEM. ns: non-significant, * $p \leq 0.05$, ** $p \leq 0.01$, *** $p \leq 0.001$.

**Suppl. Figure 6. Comparison of the transcriptome of MB G3 HDMB03 before and after the graft in the chick embryo. a.** PCA scatter plot of normalized and batch corrected gene expression comparing the pre-graft replicates (n=3) with post-graft replicates (n=3). **b.** Enrichment plots used to calculate the normalized enrichment score (NES) of "Signaling by ROBO receptors" gene set (padj and set size are indicated).

**Suppl. Figure 7. Expression of SLIT2 and its receptors in cerebellar progenitors and in MB cell lines. a.** As in Fig. 7ijk, SLIT1 and SLIT3 expression in human cerebellar progenitors derived from the RL are shown. **bcde.** The expression of ROBO1 (bd) and PLXNA1 (ce) in the proteomic (Bernardi *et al.,* under revision) and methylome[1] defined subtypes from our in-house patient samples collection, at the transcriptomic (bc) and at the protein level (de). **fg.** Average expression, at the mRNA level, of ROBO1 (f) and PLXNA1 (g) in MB tumors from WNT, SHH, G3 and G4 tumors in a public cohort of MB patient samples[2]. **hi.** Average expression, at the mRNA level, of ROBO1 (h) and PLXNA1 (i) in transcriptomic public data sets of the cell lines used in this study, except PTC1MUT which are murine cells.

**Suppl. Figure 8. SLIT signaling has no impact on HD-MB03 cells apoptosis and proliferation. ab.** Active caspase-3 (**a**) and KI67 (**b**) staining was performed on cells cultured on slices in the various conditions indicated for a period of 48h. Recombinant fragments SLIT2 N and SLIT2 C (2 mg/ml final concentration) were or were not added. Staurosporin (0,46 mg/mL final concentration) was used as a positive pro-apoptotic control. 20 % SVF medium was used as a positive pro-proliferative control. Error bars indicate SEM. The Kruskal-Wallis test was applied to compare all conditions. ns: non-significant, *** $p \leq 0.001$. **c.** Quantification of siROBO1 and siPLXNA1 impact on *ROBO1* and *PLXNA1* mRNa. The ratio between mRNA level and the housekeeping gene *GAPDH* mRNA level, i.e., the $2^{DCt}$ raw value = $2^{-(\text{Ct Gene of Interest} - \text{Ct GAPDH})}$, is shown for each condition (24 and 48h post-transfection).

**Suppl. Figure 9. SLIT signaling impacts MB G3 tumorigenesis early and is still active in patients. ab.** Kaplan-Meier overall survival analysis was performed on ROBO1 (a) and PLXNA1 (b) expression in our in-house cohort. **cd.** Kaplan-Meier overall survival analysis was performed on ROBO1 (c) and PLXNA1 (d) expression in the Cavalli cohort (Cavalli *et al,.* 2017). **ef.** Kaplan-Meier overall survival analysis was performed on $ROBO1^{high}PLXNA1^{high}$ and $ROBO1^{low}PLXNA1^{low}$ groups of MB SHH (e) and MB G4 (f) tumors.

**Suppl. Figure 10. SLIT signaling is correlated with MYC expression in MB G3.**

**ab.** ROBO1 (a) and PLXNA1 (b) expression at the transcriptomic level in the various MB G3 subtypes of the Cavalli cohort. The Kruskal-Wallis test was performed. **c.** The two groups defined by ROBO1 expression in Supp. Fig. 9a were evaluated for their expression of MYC. A T-test was performed between both groups. **d.** The two groups defined by PLXNA1 expression in Supp. Fig. 9b were evaluated for their expression of MYC. A T-test was performed between both groups. **ef.** As in Fig. 10e, Pearson correlation analysis compared MYC and ROBO1 (e) and PLXNA1 (f) expression. Coefficient of determination R2 and the pvalue P are indicated. **ghi.** The two groups $ROBO1^{high}PLXNA1^{high}$ and $ROBO1^{low}PLXNA1^{low}$ groups defined in our in-house cohort were classified based on the MB G3 methylome/transcriptomic[2] (g), proteomic (Bernardi *et al.,* under revision) (h) and methylome[1] (i) defined subtypes. **jkl.** The level of expression of ROBO1 and PLXNA1 separately is shown in the various MB G3 subtypes in our in-house cohort: methylome/transcriptomic[2] (j), proteomic (Bernardi *et al.,* under revision) (k) and methylome[1] (l). **mno.** Pearson correlation analysis comparison of MYC and ROBO1 + PLXNA1/2 (m), ROBO1 (n) and PLXNA1 (o) expression in our in-house cohort. Diamonds plot MYC amplified tumors (AMP), circules plot MYC non-amplified tumors (No_AMP) and squares indicate non defined tumors (N/A). Coefficient of determination R2 and the pvalue P are indicated. ns non-significant, * $p \leq 0.05$, ** $p \leq 0.01$, *** $p \leq 0.001$, **** $p \leq 0.0001$.

**Suppl. Figure 11. SLIT signaling favors the exploratory behavior of MB G3 cells.**

**a.** Diagram summarizing the most described interactions in the SLIT2 network. Line width is proportional to an interaction score calculated by STRINGR website. **b.** Scheme of the role we propose for SLIT signaling during early MB tumorigenesis (see text).

**Movie S1.** 3D-reconstruction of a whole HH29 (E6) chick embryo grafted with HD-MB03::GFP cells, labeled with an anti-human mitochondria antibody, cleared and imaged with SPIM.

**Movie S2.** The grafted embryo presented in Movie S1 after image treatment and surface reconstruction using Imaris software to visualize ventricles and parenchyma.

**Movie S3.** 3D-reconstruction of a whole E12 chick embryo grafted with HD-MB03::GFP cells, labeled with an anti-GFP antibody, cleared and imaged with SPIM. Transverse and sagittal optic sections are successively shown. 3D reconstruction was performed with Imaris software to better visualize the topography of the tumor. The cerebellum outlines are represented in white and the tumor mass surface reconstruction is in pink.

**Table 1. Differential RNAseq :** 2636 significantly deregulated genes (padj<0.05, absFc>1.5)) between both experimental conditions : post- versus pre-grafted HD-MB03 cells.

**Table 2. Home-made list of genes involved in axon guidance.**

**Table 3. Data associated with patients' samples.**


1. Sharma, T. *et al.* Second-generation molecular subgrouping of medulloblastoma: an international meta-analysis of Group 3 and Group 4 subtypes. *Acta Neuropathol* 138, 309–326 (2019).
2. Cavalli, F. M. G. *et al.* Intertumoral Heterogeneity within Medulloblastoma Subgroups. *Cancer Cell* 31, 737-754.e6 (2017).
