## Supplemental Table 3 for "Lineage of origin-specific developmental programs drive the behaviors of malignant cells in an avian embryo model of human Medulloblastoma subgroups"

**Table 3. Data associated with patients’ samples.**

| Age | Sample | Sex | Subgroup | MYC ampl |
| --- | --- | --- | --- | --- |
| 6 y | Biopsy | M | G3 | + |
| 2 y | Biopsy | M | SHH | - |
| 14 y | Biopsy | M | G4 | - |
